# Structure of an eDNA-scaffolded Scp-like protease filament from *Streptococcus sanguinis*

**DOI:** 10.64898/2026.05.25.726231

**Authors:** Robin Anger, Laetitia Pieulle, Ana Vallart, Lucas Baudeau, Vladimir Pelicic, Rémi Fronzes

## Abstract

C5a peptidases are surface-associated serine proteases that enable pathogenic streptococci to evade innate immunity by degrading neutrophil chemoattractants. These enzymes are usually anchored to the bacterial cell wall via an LPXTG motif and have only been characterized as monomers. Here, we report the cryo-EM structure of an unexpected supramolecular assembly from *Streptococcus sanguinis* (*S. sanguinis)*, in which a Streptococcal C5a peptidase-like (ScpH) oligomerizes into an extended filament scaffolded on extracellular DNA (eDNA). The 3.1 Å resolution structure reveals that this protease forms interlocked dimers through reciprocal contacts between the Ig and protease-Fn domains, and that these dimers propagate into a double-rowed filament via three additional inter-protomer interfaces. The central eDNA strand is engaged by an electropositive cleft formed by the C-terminal FIVAR3 domain and the DNA-binding tail, which replace the LPXTG anchor found in characterized orthologs. AlphaFold 3 predictions indicate that both dimerization and eDNA binding are conserved across a subfamily of streptococcal species that are prominent oral colonizers and eDNA producers. The catalytic triad is structurally intact and a co-purified peptide occupies the active site, although proteolytic activity remains to be demonstrated. This structure reveals a previously unknown mechanism of biofilm functionalization in which a putative immune evasion protease is assembled directly onto eDNA, transforming a passive structural component of the biofilm matrix into a potential catalytic platform.

## Introduction

As a prominent member of the human oral microbiota, *Streptococcus sanguinis* is a primary colonizer of the tooth surface and a major component of dental plaque biofilm^1,2^. Although primarily a beneficial commensal that antagonizes cariogenic pathogens such as *Streptococcus mutans*^3,4^, it is also a notorious opportunistic pathogen when it enters the bloodstream^5^. Transient bacteremia, often triggered by routine dental procedures or inadequate oral hygiene, allows *S. sanguinis* to reach and adhere to damaged cardiac endothelium, making it an important cause of infective endocarditis^6–8^.

To survive and establish an infection in this demanding environment, where it faces constant surveillance and rapid attacks by infiltrating neutrophils^9^, antimicrobial platelets^10^, and the complement system^11^, *S. sanguinis* must successfully overcome the host’s early innate immune defenses. It achieves this through a multifaceted evasion strategy: inducing neutrophil cell death via the production of hydrogen peroxide (H_2_O_2_)^12^, escaping neutrophil extracellular traps (NETs) through the activity of a cell wall-anchored nuclease^13^, assembling functional amyloid fibrils to significantly reduce its susceptibility to complement-mediated clearance^11^ and hijacking platelets via surface adhesins to form protective vegetations that shield it from immune clearance^14^.

In related pathogenic species, such as *Streptococcus pyogenes* (Group A *Streptococcus*, GAS) and *Streptococcus agalactiae* (Group B *Streptococcus*, GBS), a key mechanism of immune evasion involves the expression of surface-associated serine proteases belonging to the S8 (subtilisin-like) family, which specifically cleave neutrophil chemoattractants^15^. By degrading these crucial signaling molecules, these pathogens effectively abrogate the recruitment of neutrophils to the site of infection^15^. Notable examples of such serine proteases include the peptidases ScpA (in *S. pyogenes)* and ScpB (in *S. agalactiae*), as well as the protease ScpC (also known as SpyCEP in *S. pyogenes*)^16–18^. These enzymes exhibit a highly conserved structural organization: they are canonically anchored to the bacterial cell wall via a C-terminal LPXTG motif and possess a central catalytic domain built around a strictly conserved aspartate-histidine-serine triad^16–18^.

Despite their structural similarities, these proteases exhibit distinct substrate specificities, targeting different components of the innate immune response. The peptidases ScpA and ScpB both act as anaphylatoxin inactivators, but they differ significantly in their substrate specificity. While ScpB is currently known to strictly target the C5a anaphylatoxin^17^, the substrate repertoire of ScpA is broader, encompassing C5a as well as other complement factors (C5, C3 and C3a) and pro-inflammatory cytokines, such as IL-37, type II (IFN-γ) and type III (IFN-λ1 and IFN-λ2) interferons^19,20^. The neutralization of the C5a anaphylatoxin, for example, occurs through a highly specific proteolytic cleavage between His67 and Lys68, removing the C-terminal region essential for receptor binding^21^. In contrast, the protease ScpC (also known as SpyCEP) specializes in the inactivation of chemokines such as CXCL1, CXCL2, CXCL3, CXCL5, CXCL6, CXCL7 and CXCL8^22–24^. For example, ScpC cleaves the CXCL8 C-terminus between Gln59 and Arg60, thereby preventing this chemokine from binding to the CXCR1 and CXCR2 receptors on neutrophils^25^.

A genomic analysis of *S. sanguinis* has revealed that it possesses a homologous gene encoding a subtilisin-like serine protease closely related to the above C5a peptidases^15,26,27^. However, unlike its well-characterized orthologs, sequence analysis indicates that this *S. sanguinis* homolog completely lacks the canonical LPXTG cell wall-anchoring motif. Consistent with this atypical organization a recent study highlighted that this putative enzyme is highly abundant in *S. sanguinis* extracellular membrane vesicles (EMVs)^28^. Crucially, deletion of the corresponding gene not only significantly increases the internalization of these EMVs by gingival epithelial cells, but also abolishes their ability to inhibit host interferon signaling, which suggested that the enzyme normally suppresses this early immune response, potentially by directly cleaving host interferons^29^. While the precise physiological consequences of this EMV-associated activity are still emerging, the strong homology of this gene to the chemoattractant-cleaving proteases of GAS and GBS suggests that this enzyme could serve as an additional weapon in the rich immune evasion arsenal of *S. sanguinis*. Yet, while the three-dimensional architecture and structural basis of substrate recognition are well established for ScpA, ScpB, and ScpC in pathogenic streptococci, the structural organization and activity of the *S. sanguinis* homolog remains unknown.

Virulence-associated functions are not limited to the bacterial cell surface but can also be deployed within the extracellular matrix of biofilms. This matrix is a complex assembly of exopolysaccharides, proteins, lipids, and extracellular DNA (eDNA), which together provide structural scaffolding, mediate intercellular cohesion, and shield resident bacteria from host immune defenses^30,31^. Among these components, eDNA has emerged as a particularly critical structural element, as its enzymatic degradation destabilizes early-stage biofilms^31^. Importantly, growing evidence indicates that the biofilm matrix is not merely a passive scaffold or a shield but can also harbor catalytic functions. In *Pseudomonas aeruginosa*, biofilms have been shown to harbor enzymatically active proteins that can be retained through interactions with structural exopolysaccharides, where they contribute to immune defense and nutrient acquisition^32–34^.

Here, we present the high-resolution cryo-EM structure of a previously unrecognized supramolecular assembly from *S. sanguinis*, in which a subtilisin-like serine protease homologous to the streptococcal C5a peptidases is assembled onto extracellular double-stranded DNA (eDNA). We demonstrate that, rather than being covalently tethered to the cell wall like its well-characterized orthologs in other pathogenic streptococci, this secreted protease binds to eDNA via a lysine-rich C-terminal tail. This interaction yields a continuous nucleoprotein filament, turning bare eDNA into a functionalized, protease-decorated scaffold. This discovery redefines Scp-like proteases from simple cell-surface enzymes to components of a higher-order virulence architecture, revealing a sophisticated mechanism of biofilm functionalization.

## Results

### Serendipitous discovery of an unknown filamentous assembly in *S. sanguinis* 2908 sheared fractions

In a previous study, we determined the cryo-EM structure of the type 4 pilus (T4P) from *S. sanguinis* using material obtained by mechanical shearing of surface-associated filaments^35^. During that work, two major types of filaments were identified on the cryo-EM grids: the ∼7 nm-wide T4P that were the focus of the study, and thin ∼3 nm-wide filaments consistent with naked eDNA^35^. Upon re-examination of the data after the study was published, we noticed the presence of a third, previously overlooked species displaying a larger diameter and a strikingly repetitive arrangement (Fig. 1a). 2D class averages revealed an ordered assembly with a clear internal periodicity, distinct from both T4P and bare eDNA (Fig. 1b).

**Figure 1:**
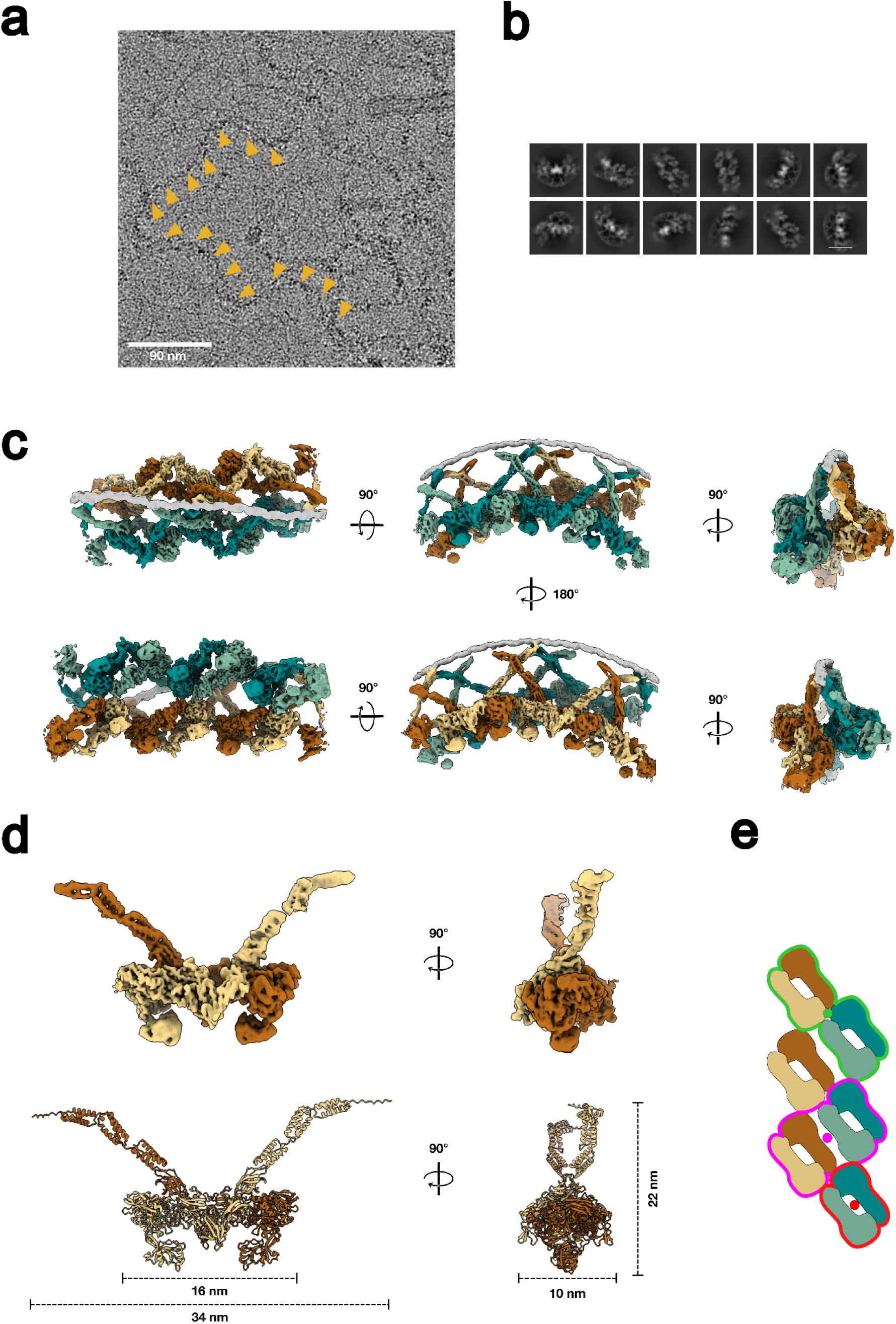
Cryo-EM structure and overall architecture of the ScpH filamentous assembly. **(a)** Representative cryo-EM micrograph of *S. sanguinis* sheared surface extracts. Yellow arrowheads highlight the repetitive ScpH filamentous assemblies. Scale bar: 90 nm. **(b)** 2D classes averages of ScpH filaments. Scale bar: 190 Å. **(c)** Consensus cryo-EM map at 3.2 Å resolution. The architecture resembles a suspension bridge, with two parallel rows of ScpH protomers flanking a central filament of unknown nature (grey). The two rows are colored in shades of green and brown, respectively, with individual protomers within each row distinguished by alternating light and dark hues (light/dark green and light/dark brown). **(d)** Cryo-EM map and ribbon representation of orthogonal views of an interlocked ScpH homodimer. The repeating building block spans approximately 34 nm in length, 10 nm in width, and 22 nm in height. **(e)** Schematic diagram of the filament assembly illustrating the protomer packing and the network of C2 symmetry axes. The different symmetry relationships are outlined in red, pink, and green, with each corresponding C2 axis symbolized by a colored dot.

To identify the molecular components of this assembly, we analyzed the sheared fraction by SDS-PAGE followed by mass spectrometry. Among the major protein species detected, we identified a subtilisin-like serine protease (locus tag SSV_1730 in *S. sanguinis* strain 2908, corresponding to SSA_1882 in the reference strain SK36) homologous to the streptococcal C5a peptidases ScpA and ScpB (Fig. S1a). We hereafter refer to this protein as ScpH (Streptococcal C5a peptidase group H)^36^. To obtain a high-resolution structure, we then performed a dedicated purification to enrich these assemblies and collected a new cryo-EM dataset, which forms the basis of the structure presented below.

### Cryo-EM reconstruction of the ScpH filamentous assembly

To determine the structure of the ScpH assembly at high resolution, we developed a dedicated purification protocol using *S. sanguinis* SS56 strain in which the *pil* locus encoding T4P was entirely deleted. Bacteria were grown in liquid culture, and surface-associated material was released by controlled mechanical shearing of the cell pellets. After clearing of intact cells and debris, the filaments were pelleted by ultracentrifugation at 100,000 × g and resuspended in a minimal volume for cryo-EM grid preparation.

Cryo-EM data were collected on a Glacios 2 microscope operated at 200 kV, equipped with a Falcon 4i detector and a SelectrisX energy filter, at a pixel size of 0.885 Å. A total of 25,079 movie stacks were collected, of which 22,037 micrographs were retained after motion correction and CTF estimation. Particle picking was performed using Topaz through an iterative training loop, yielding 202,815 initial particles. After successive rounds of *ab initio* reconstruction and heterogeneous refinement in cryoSPARC, a cleaned subset of 65,221 particles was used to generate a consensus reconstruction at 3.2 Å resolution with a C2 symmetry imposed (Fig. 1c, Fig. S2a and Fig.S3).

The resulting map reveals a striking architecture in which two parallel rows of ScpH protomers run side by side beneath a central filament of unknown nature (Fig. 1c). Each protomer is connected to the central filament through a long arch, while the remaining domains form a continuous deck through extensive contacts both within each row along the filament axis and between the two rows in a lateral fashion. The overall arrangement is reminiscent of a suspension bridge, in which the deck is formed by the interlocking protein domains while the central filament runs along the top like a supporting cable (Fig. 1c).

The repeating building block of the assembly is a dimer of ScpH protomers related by a local two-fold (C2) symmetry axis (Fig. 1d). Each dimer spans approximately 34 nm in length, 10 nm in width, and 22 nm in height (Fig. 1d). At the level of the filament, two additional C2 symmetry axes relate the two rows to one another. A schematic representation of the protomer packing and symmetry relationships is shown in Fig. 1e.

### Structure of the ScpH monomer and domain organization

The ScpH monomer spans 1,506 residues and adopts an elongated, multi-domain architecture. Sequence analysis combined with AlphaFold 3 prediction of the full-length monomer reveals the following domain organization (Fig. 2a,b): a pre/pro region (residues 1–173), a protease domain with an inserted PA subdomain (protease: 174–411 and 556–670; PA: 412–555), three fibronectin type III (Fn) domains (Fn1: 671–798; Fn2: 799–994; Fn3: 995–1105), two immunoglobulin-like (Ig) domains (Ig1: 1106–1196; Ig2: 1197–1281), three FIVAR domains (FIVAR 1: 1282–1347; FIVAR 2: 1348–1419; FIVAR 3: 1420–1492), and a C-terminal domain of unknown function, here termed Not Determined (ND) (1493–1506) (Fig. 2b). The protease domain harbors a strictly conserved catalytic triad (Asp–His–Ser) characteristic of S8 subtilisin-like serine proteases (Fig. 2b). The PA subdomain, inserted within the protease domain, is a hallmark of this family and has been implicated in substrate recognition in related enzymes. Within the filament, the protease, PA, Fn, and Ig domains form the deck of the assembly, while the FIVAR and ND domains constitute the elongated arch connecting each protomer to the central filament.

**Figure 2:**
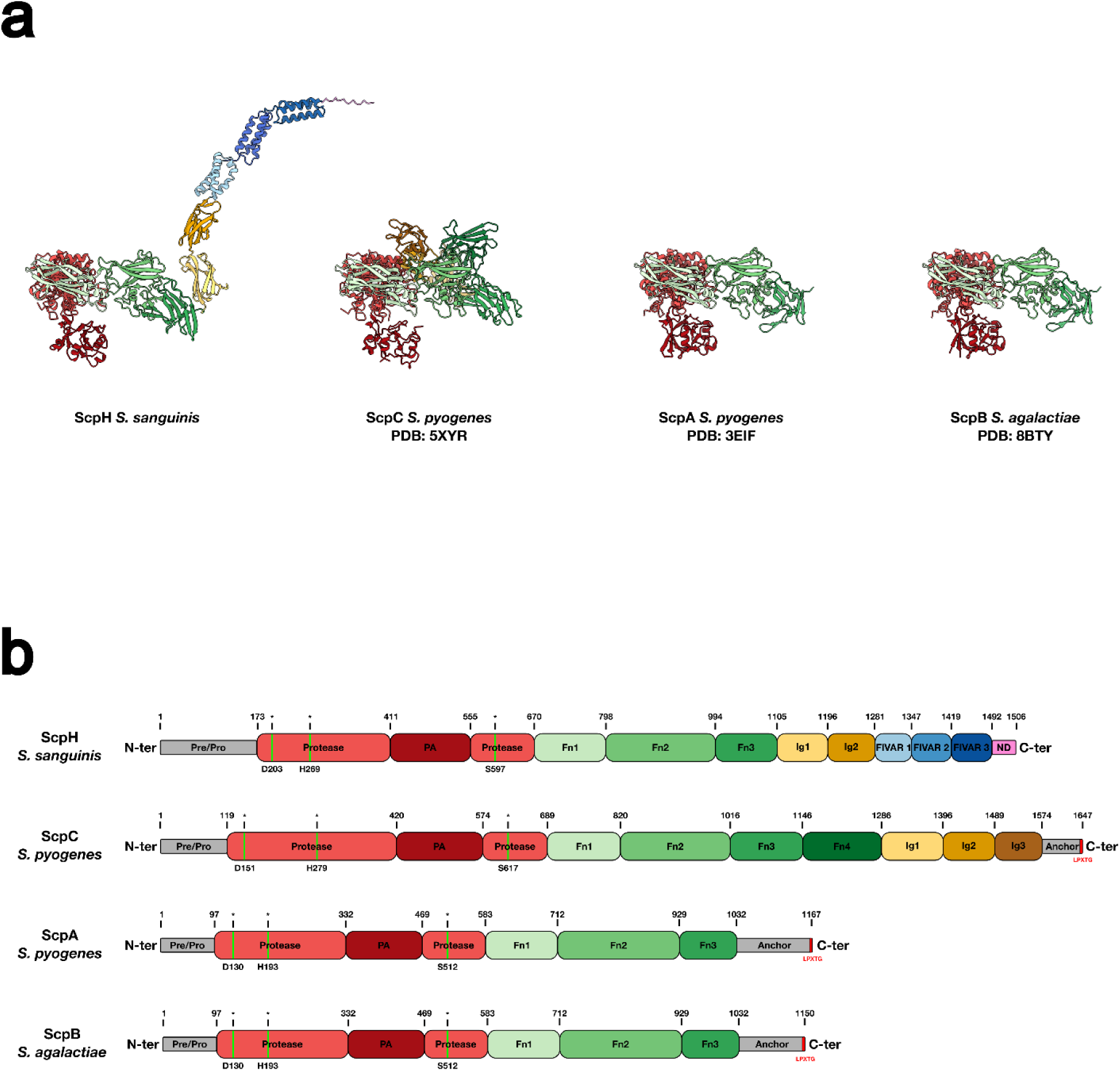
Domain organization of ScpH and structural comparison with streptococcal orthologs. **(a)** Ribbon representation of ScpH monomer of *S. sanguinis* (residues 173-1506, AlphaFold 3 predicted fold), ScpC (residues 119-1574) of *S. pyogenes* (PDB: 5XYR), ScpA (residues 97-1032) of *S. pyogenes* (PDB: 3EIF) and ScpB (residues 97-1032) of *S. agalactiae* (PDB: 8BTY). **(b)** Linear schematic representation of the domain organization of the various Scp proteases, where the amino acids forming the catalytic triad are indicated within the protease domain by a green vertical line and an asterisk. **(a, b)** The color code for the different domains is consistent across both panels. The protease domain is shown in red, the PA subdomain in dark red, the fibronectin type III (Fn) domains in shades of green, and the immunoglobulin-like (Ig) domains in shades of yellow. The FIVAR domains, shown in shades of blue, and the C-terminal Not Determined (ND) domain, shown in pink, are exclusive to ScpH. The N-terminal Pre/Pro regions of the different Scp proteins and the C-terminal cell wall anchors of the orthologs are indicated in grey in (b) but are not represented in the structural models in (a). Finally, the canonical LPXTG motif within the anchors of the orthologs is indicated in red.

Structural comparison with the characterized orthologs ScpA from *S. pyogenes* (PDB: 3EIF), ScpC from *S. pyogenes* (PDB: 5XYR), and ScpB from *S. agalactiae* (PDB: 8BTY) reveals that the N-terminal protease core, PA subdomain, and Fn domains are conserved across the family (Fig. 2a,b). However, *S. sanguinis* ScpH differs markedly from its orthologs in the C-terminal region. ScpA (*S. pyogenes*, 1167 residues) and ScpB (*S. agalactiae*, 1150 residues) both terminate after Fn3 with a canonical LPXTG cell wall-anchoring motif. ScpC (*S. pyogenes*, 1647 residues), the largest family member, extends further with an Fn4 domain and three Ig-like domains but retains an LPXTG anchor. In contrast, *S. sanguinis* ScpH (1506 residues) replaces these C-terminal features with two Ig domains, three FIVAR domains, and a terminal ND domain, with no LPXTG motif (Fig. 2b). This absence of a canonical cell wall anchor distinguishes ScpH from all characterized orthologs and raises the question of how this protease is presented at the bacterial surface, a point that is directly addressed by the supramolecular assembly described here.

To obtain a high-resolution experimental model, we exploited the global C2 symmetry relating the two rows of the filament by performing symmetry expansion, generating 130,442 pseudo-particles that were subjected to focused local refinement on the deck region of the complex (see Methods and Fig. S3). The refined volume encompasses the asymmetric unit at the base of the assembly: two complete dimers spanning both rows, together with an Fn1 part of the adjacent dimer within the same row, thereby capturing all four inter-protomer interfaces described below. This yielded a map at 3.1 Å resolution in which the protease, Fn, and Ig domains were resolved at sufficient quality for *de novo* model building (Fig. S2b and S4a). The atomic model was built and refined against this locally refined map, providing an experimentally determined structural framework for all inter-protomer contacts in the assembly (Fig. S4b).

To complete the structure, we supplemented the experimentally determined regions with AlphaFold 3 predictions. The PA subdomain and the arch regions (FIVAR 1–3 and ND), poorly resolved in the experimental map, were derived from the AlphaFold 3 prediction of the full-length monomer and rigid-body docked into the consensus cryo-EM density. The resulting composite model and the boundaries between experimentally determined and predicted regions are indicated in Fig. S4a.

### Inter-protomer interfaces drive oligomerization

The repetitive arrangement of ScpH protomers within the filament is mediated by an extensive network of inter-protomer contacts. Four distinct interfaces, labeled I–IV, were identified in the structure (Fig. 3).

**Figure 3:**
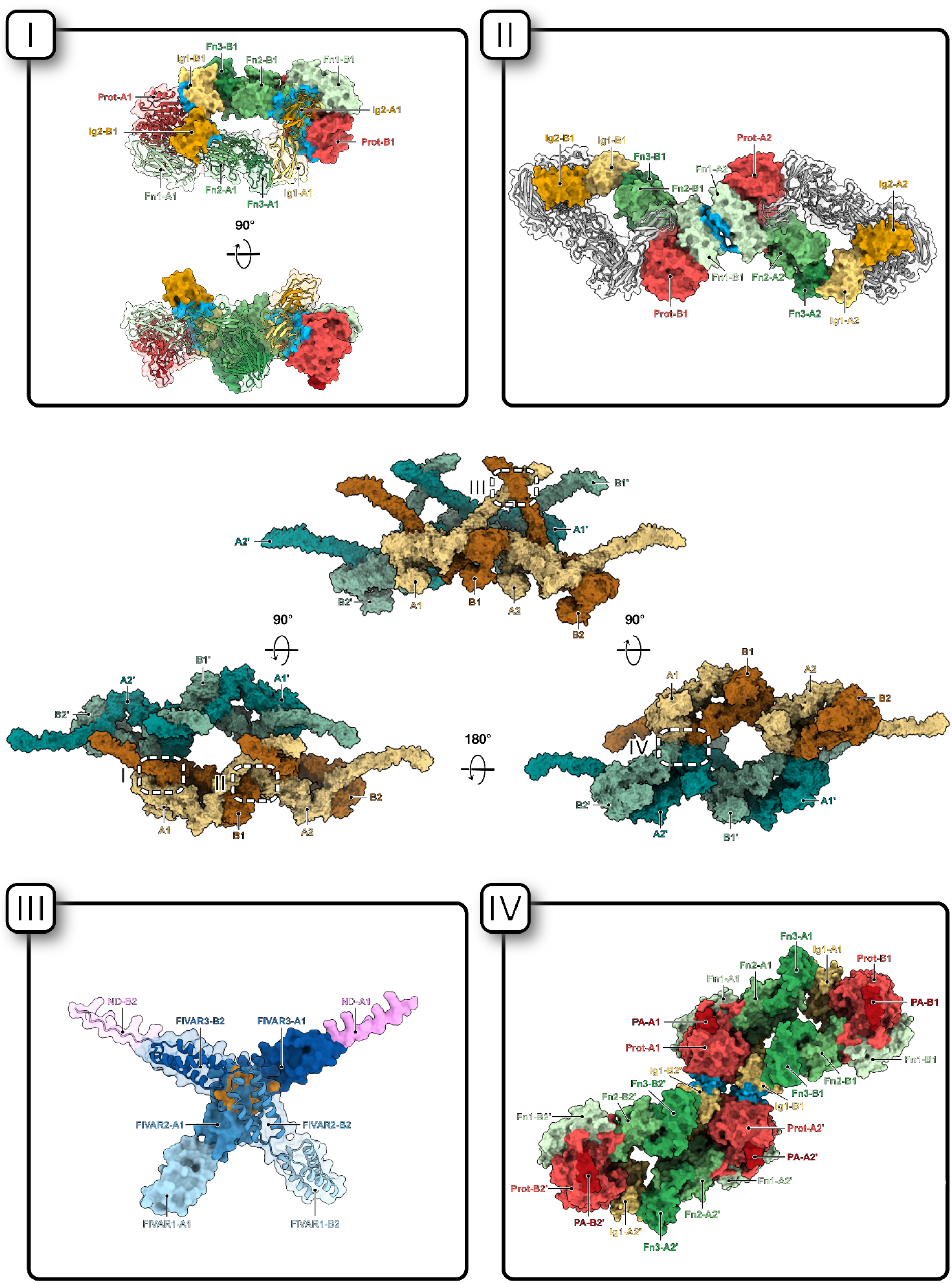
Structural basis of ScpH oligomerization and inter-protomer interfaces within the filament. The central panels display a surface representation of the overall architecture of the ScpH filament assembly from multiple angles (top, side, and bottom views), using the same color code as in Figure 1c. This illustrates the arrangement of individual protomers that constitute the double-rowed structure, where protomers in one row are labeled A1, B1, A2, B2 and those in the opposite row are labeled A1’, B1’, A2’, B2’. The four distinct inter-protomer interfaces (I-IV) driving this supramolecular assembly are boxed and detailed in the surrounding expanded panels, where the domain color coding is consistent with Figure 2. For each interface, the contact zone is highlighted in blue for interfaces I, II, and IV, and in orange for interface III. **(I)** Interface I represents the primary intra-dimer contact that stabilizes the fundamental repeating building block of the assembly, forming a closed elliptical ring involving the Ig1 and Ig2 domains of one subunit interacting with the protease, Fn1, and Fn2 domains of its partner subunit. **(II)** Interface II connects successive dimers longitudinally within the same row of the filament deck and is formed directly between the Fn1 domains of adjacent dimers. **(III)** Interface III further stabilizes the longitudinal connection between successive dimers at the level of the protruding arches through contacts between the FIVAR2 domains of adjacent dimers. **(IV)** Interface IV constitutes the lateral inter-row contact bridging the two parallel rows together through specific contacts between the protease domain of a dimer in one row and the Ig1 domain of a dimer in the adjacent row.

Interface I is an intra-dimer contact that holds together the two subunits forming the repeating building block. It is formed between the Ig1 and Ig2 domains of one subunit, and the protease, Fn1 and Fn2 domains of the partner subunit. Owing to the local C2 symmetry within the dimer, this interface is duplicated: each subunit reciprocally contributes its Ig1–Ig2 domains to contact the protease, Fn1 and Fn2 domains of its partner. Together, these reciprocal contacts create a closed elliptical ring in which the two protomers are interlocked (Fig. 3, panel I), providing a robust structural foundation for each dimer upon which the elongated arches extend upward toward the central filament. This is the most extensive interface in the assembly, burying ∼2,550 Å² of surface area per contact. The Ig1 domain engages primarily the protease domain through an extensive network of polar contacts, including a salt bridge (Lys1132–Asp239) and hydrogen bonds involving Arg1130, Tyr1138, Lys1143, and Asp1165 on the Ig1 side and Tyr252, Asp256, and Ser350 on the protease side. The Ig2 domain bridges three domains: it contacts the protease through Arg1238 and Ser1239 (engaging Asp346, Val347, and Gln352), the Fn2 domain through Asn1271 (engaging Asn906 and Thr976), and the Fn1 domain through additional buried contacts. In total, PISA analysis identifies 16 hydrogen bonds and one salt bridge across this interface. This region of the map is well resolved, lending confidence to the identification of the interacting residues.

Interfaces II and III connect successive dimers within the same row. Interface II is formed between the Fn1 domains of adjacent dimers, burying ∼440 Å² of surface area (Fig. 3, panel II). Despite its modest size, this contact creates a direct link between the deck regions of neighboring dimers, ensuring longitudinal continuity along each row. Interface III is located at the level of the arches. As can be seen in the isolated dimer structure (Fig. 1d), the arches extend beyond the boundaries of the dimer, reaching out toward the neighboring n+1 and n−1 dimers within the row. These protruding arches are connected through interactions between the FIVAR 2 domains of adjacent dimers, with a contribution from FIVAR 3, burying ∼860 Å² of surface area (Fig. 3, panel III). PISA analysis of the AlphaFold 3-predicted FIVAR dimer indicates that the interface is predominantly electrostatic in nature.

Interface IV bridges the two rows, connecting dimers from opposite sides of the filament through limited contact (∼210 Å² buried surface area) between the protease domain of one dimer and the Ig1 domain of a dimer in the adjacent row (Fig. 3, panel IV). Due to the global C2 symmetry of the filament, this interface is duplicated, generating symmetric pairs of inter-row contacts.

Although interfaces II, III, and IV are individually modest in size, they are all repeated along the entire length of the filament. Their additive nature likely contributes significantly to the overall stability of the assembly, ensuring both the longitudinal propagation of each row (interfaces II and III) and the lateral cohesion between the two rows (interface IV), complementing the bridging role of the central filament.

Notably, the Ig1 domain emerges as a keystone of the assembly: it participates both in the intra-dimer interface I and in the inter-row interface IV, making it a central hub for the structural integrity of the filament. For interfaces II, III, and IV, where the local resolution is more limited or the model derives from predictions, the buried residues are highlighted in the zoom panels of Fig. 3 but specific contacts should be interpreted with caution.

### The central filament is double-stranded DNA

Having established the overall architecture and inter-protomer contacts of the ScpH assembly, we next sought to determine the nature of the central filament running along the apex of the structure. A focused local refinement was performed on this region using the consensus particle set (see Methods and Fig.S3), yielding a map in which the central filament was resolved with sufficient detail to reveal a characteristic helical pitch and major/minor groove pattern consistent with B-form double-stranded DNA (Fig. 4a, Fig.S2c).

**Figure 4:**
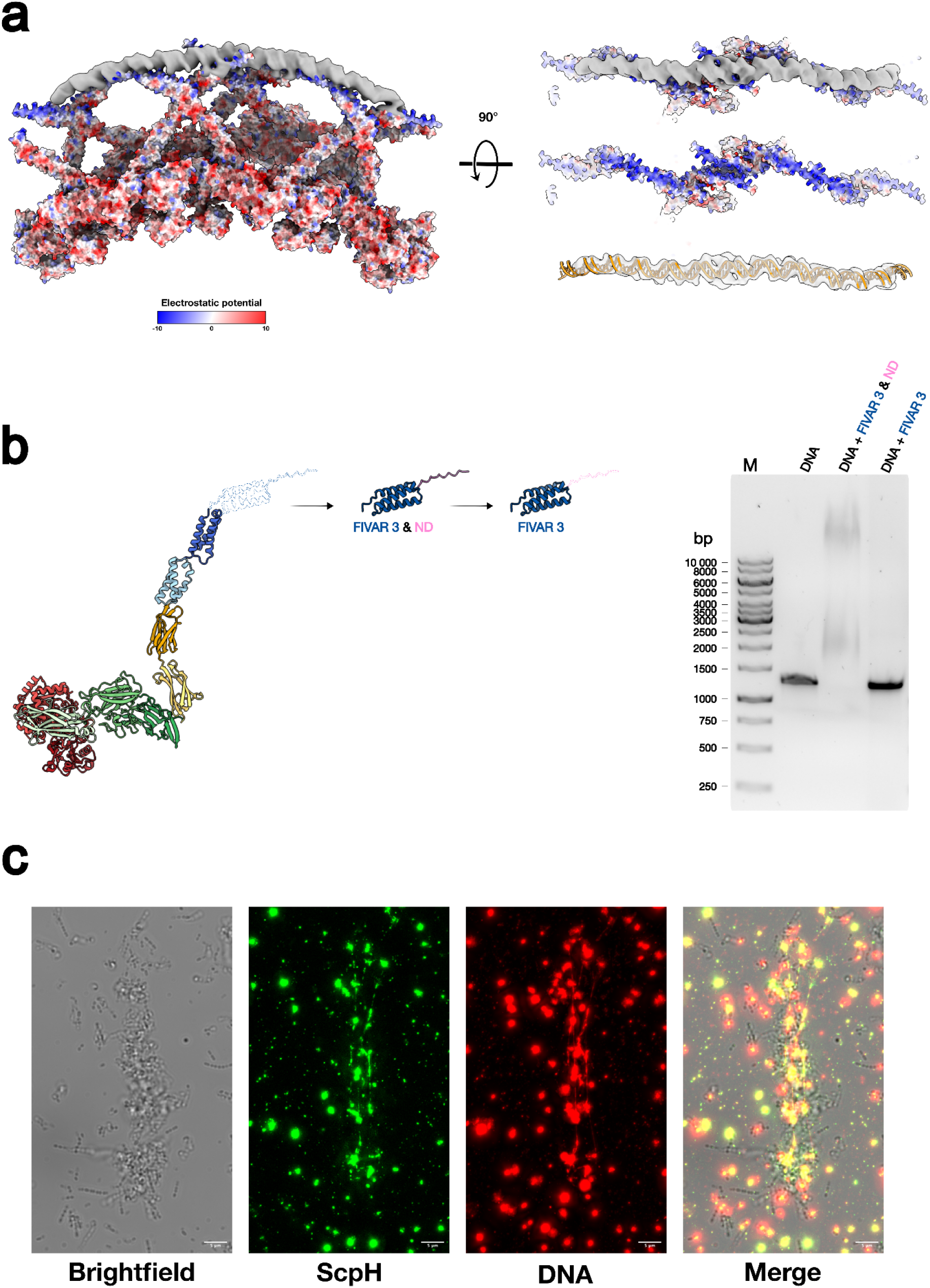
Structural and functional basis of the ScpH interaction with eDNA. **(a) Visualization of the central filament and electrostatic surface potential.** Left: Surface representation of the ScpH filament colored by electrostatic potential (red, negatively charged and blue, positively charged), with the locally refined map of the central density shown in grey. Right: 90° rotated views of the C-terminal region model (FIVAR 3 and ND domains) highlighting a strongly electropositive cradle. This positively charged surface faces the central filament, which displays the characteristic helical pitch and major/minor groove pattern of B-form double-stranded DNA. **(b) *In vitro* DNA-binding by the ScpH C-terminal tail.** Left: Ribbon diagrams of the recombinant constructs (FIVAR 3 & ND, and FIVAR 3 alone). Right: Electrophoretic mobility shift assay (EMSA) of a 1,297 bp DNA fragment incubated with the indicated proteins. The FIVAR 3 & ND construct induces a clear gel shift, whereas the FIVAR 3 domain alone does not, demonstrating that the terminal ND domain is functionally essential for stable DNA binding. **>(c) *In vivo* co-localization of ScpH and eDNA.** Immunofluorescence microscopy of *S. sanguinis* cultures. Panels display bacterial cells (Brightfield), the ScpH protease detected via an anti-ALFA nanobody (green) and eDNA labeled with an anti-DNA antibody (red). The merged image confirms the co-localization of ScpH and DNA signals around the cells. Scale bars: 5 µm.

This identification prompted us to examine how ScpH interacts with DNA. Mapping of the electrostatic surface potential onto the AlphaFold 3-predicted model of the C-terminal region revealed that the FIVAR 3 domain and the ND domain display a strongly electropositive surface (Fig. 4a). In the context of the assembly, these positively charged patches face the DNA directly, providing a straightforward structural rationale for the interaction through charge complementarity with the negatively charged phosphate backbone in DNA.

To further investigate the interaction with DNA, we performed an AlphaFold 3 prediction of two copies of the FIVAR 1–3 and ND domains together with a 45 bp random double-stranded DNA sequence. The resulting model could be readily fitted into the cryo-EM density map (Fig. S5). Importantly, the predicted protein dimer was essentially identical to the model obtained without DNA (mean pLDDT ∼85.5 vs ∼89.5; FIVAR 2–FIVAR 2 PAE = 3.8 vs 2.8), indicating that the dimerization interface is not dependent on the presence of DNA (Fig. S5). Rather, the FIVAR 2-mediated dimer appears to form an autonomous structural unit that creates a cleft for DNA binding. The model confirms the importance of the electropositive patch on FIVAR 3 for DNA binding. The ND domain, however, was predicted with lower confidence in its positioning relative to the DNA (mean PAE = 27.5), leaving its precise contribution to the interaction unresolved by computational approaches alone.

To address this point experimentally, we produced and purified recombinant proteins encompassing the C-terminal region of ScpH and performed electrophoretic mobility shift assays (EMSA). Incubation of a 1,297 bp DNA fragment with the recombinant purified FIVAR 3 & ND protein resulted in a clear shift of the DNA band, confirming direct binding (Fig. 4b). In contrast, the purified recombinant FIVAR 3 domain alone did not induce a detectable shift (Fig. 4b). Thus, despite its low prediction confidence, the ND domain is functionally essential for stable DNA binding. In light of these findings, we hereafter refer to this C-terminal domain as the DNA-binding tail.

Finally, to assess whether ScpH and eDNA co-localize in a physiological context, we constructed a *S. sanguinis* strain producing a ScpH-ALFA-tag fusion and performed immunofluorescence microscopy on *S. sanguinis* cultures using an anti-ALFA-tag nanobody to detect ScpH and an anti-DNA antibody to label eDNA. The merged images show co-localization of ScpH and eDNA signals around bacterial cells (Fig. 4c), consistent with the formation of ScpH-eDNA assemblies in the extracellular environment.

### The predicted catalytic site in ScpH cryoEM map is occupied by a co-purified peptide

The protease domain of ScpH harbors a catalytic triad composed of Asp–His–Ser, which is clearly resolved in the cryo-EM density (Fig. 5a). The spatial arrangement of these three residues is fully consistent with the geometry expected for an active S8 subtilisin-like serine protease.

**Figure 5:**
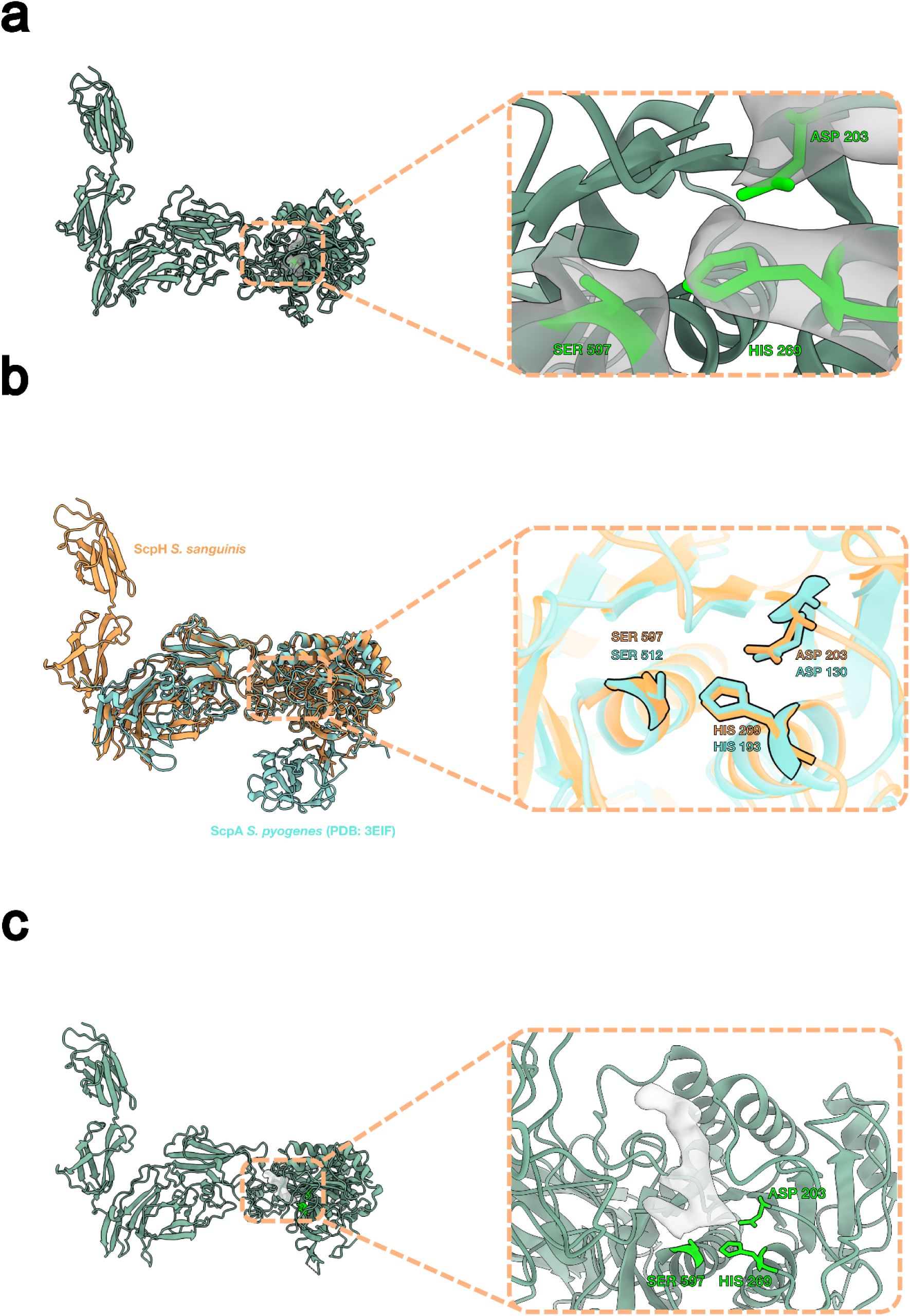
Structural conservation of the ScpH active site and identification of a co-purified peptide. **(a)** Cryo-EM density of the catalytic triad. Left: Ribbon representation of the ScpH monomer. Right: Zoomed-in view of the active site showing the strictly conserved Asp203, His269, and Ser597 residues clearly resolved within the experimental map. **(b)** Structural superposition of the ScpH catalytic triad (orange) with that of the *S. pyogenes* ortholog ScpA (light blue, PDB: 3EIF). The local structural alignment reveals strict geometric conservation of the catalytic machinery, with a root-mean-square deviation (RMSD) of 0.24 Å over the 3 Cα atoms. **(c)** Unmodeled density in the active site cleft of ScpH. The cryo-EM map reveals an additional density (grey surface) occupying the substrate-binding groove in close proximity to the catalytic triad, consistent with a short, co-purified peptide.

To characterize the structural conservation of ScpH, we focused on its N-terminal core (residues 176–1280), which encompasses the domains homologous to characterized members of the C5a peptidase family. Structural superposition of this region with *S. pyogenes* ScpA (PDB: 3EIF) reveals a high degree of overall structural conservation, with a global RMSD of 1.08 Å over 451 pruned Cα atoms. Given that relative domain flexibility can artificially inflate the global RMSD, we performed a local structural alignment restricted to the catalytic triad residues (Asp–His–Ser) to precisely assess the integrity of the active site. This targeted alignment confirmed a strict geometric conservation of the catalytic machinery, with a local RMSD of 0.24 Å over the 3 Cα atoms (Fig. 5b).

Strikingly, closer inspection of the cryo-EM map revealed an additional, unmodeled density occupying the active site cleft (Fig. 5c). This density is consistent with a short peptide and likely corresponds to a co-purified by-product of proteolytic cleavage of a protein present in the culture medium. Its position within the substrate-binding groove is compatible with the mode of substrate engagement described for the *S. pyogenes* orthologs (Fig. 5b).

To further assess whether the ScpH active site is compatible with C5a recognition, we performed an AlphaFold 3 co-fold prediction of the ScpH dimer in a complex with two human C5a (Fig. S5 and Fig. S6a). In the resulting model, C5a is positioned within the substrate-binding groove of the protease domain, with the C5a known cleavage site (His67-Lys68) located in close proximity to the catalytic triad (Fig. S6a). This prediction suggests that the active site architecture of ScpH is structurally compatible with C5a engagement, which is consistent with the conservation of the catalytic machinery observed across the family. Additionally, superposition of this model with our experimental cryo-EM map reveals that the predicted C5a cleavage site passes directly through the unassigned density found in the active site (Fig. S6b). While no definitive functional conclusions can be drawn from this *in silico* prediction alone, this spatial coincidence illustrates that the experimental density occupies the expected substrate-binding trajectory.

Despite the structural evidence for a competent catalytic site, *in vitro* enzymatic assays failed to demonstrate C5a cleavage. Following established protocols for characterizing streptococcal C5a peptidases^19^, human complement component C5a was incubated with either *S. sanguinis* whole cells or purified ScpH filaments, and the mixtures were subsequently analyzed by SDS-PAGE and Western blotting. However, these assays did not yield detectable proteolytic activity under the conditions tested. Consequently, the lack of cleavage is likely due to either a highly stringent substrate specificity, suggesting human C5a may not be the true physiological target for ScpH, or the active site being sterically occluded by the strongly bound co-purified peptide observed in our cryo-EM map. Taken together, these observations establish ScpH as a structurally competent putative peptidase whose catalytic properties remain to be fully characterized.

### Global architecture of the ScpH-DNA filament

To visualize the global organization of the filament, we performed an additional reconstruction using a larger box size encompassing up to five consecutive dimers (Fig. 6). The processing workflow and map quality are presented in Supplementary Figures. S2d and S3. This reconstruction reveals that the filament is not a straight, planar assembly but adopts a gently twisted ribbon-like morphology, in which the flat deck of the filament progressively rotates along the longitudinal axis. Whether this twist follows a regular periodicity could not be determined from the present data.

**Figure 6:**
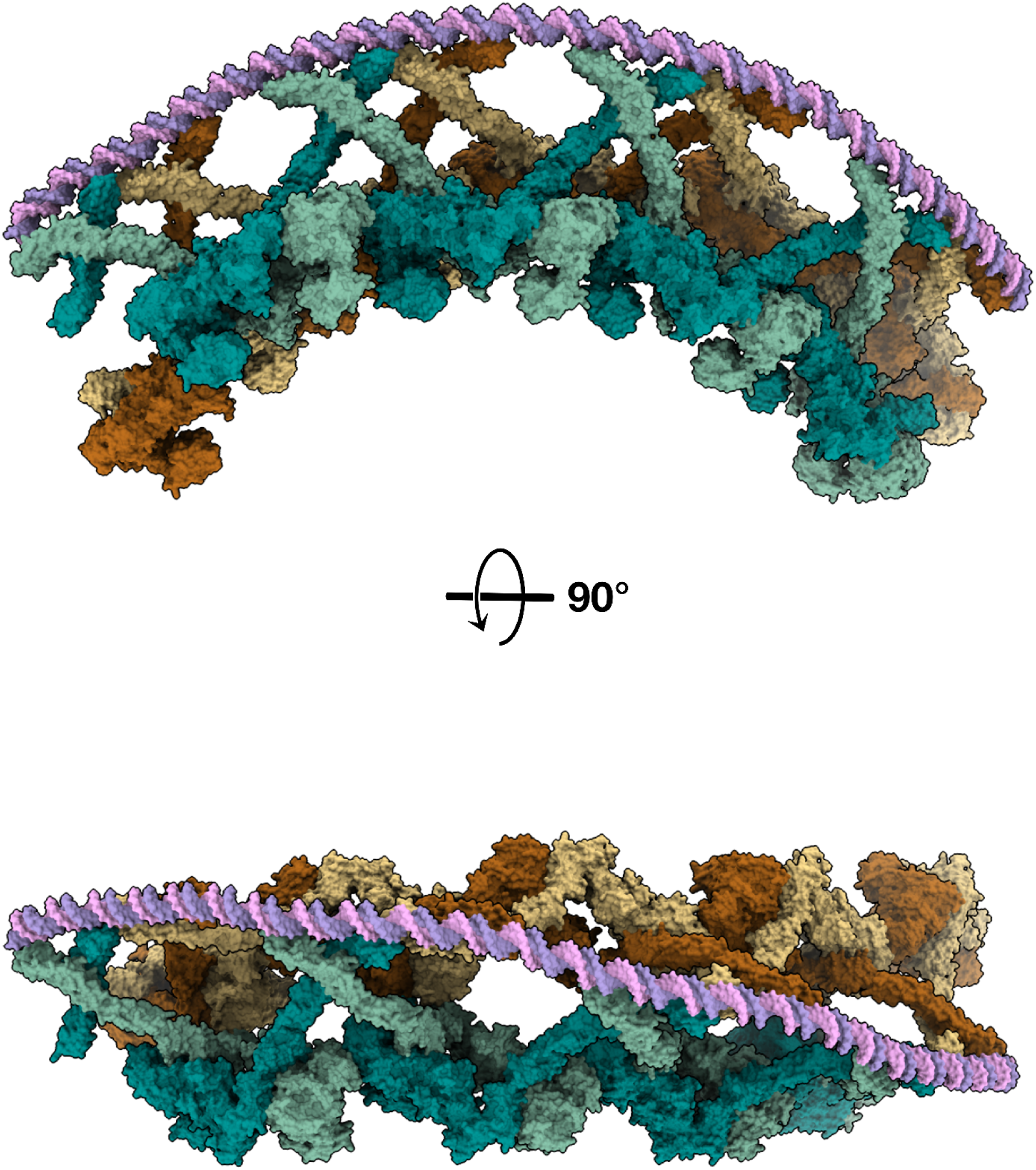
Global architecture of the extended ScpH filament. Cryo-EM reconstruction of the ScpH filament using a larger box size encompassing up to five consecutive dimers. ScpH protomers are colored according to the color code in Figure 1c, and the two DNA strands are highlighted in pink and purple. Orthogonal views (rotated by 90°) reveal that the supramolecular assembly is not a straight, planar structure, but instead adopts a gently twisted ribbon-like morphology.

## Discussion

The cryo-EM structure of the ScpH filament reveals several novel features of C5a peptidase biology that redefine our understanding of how these immune evasion factors can be organized at the bacterial surface.

First, the ScpH dimer, stabilized by the extensive Interface I between the Ig1-Ig2 and protease-Fn1-Fn2 domains, represents to our knowledge the first observation of C5a peptidase dimerization. All previously characterized family members, ScpA from *S. pyogenes*, ScpB from *S. agalactiae*, and ScpC from *S. pyogenes*, have been described exclusively as monomers, and AlphaFold 3 co-fold predictions of these orthologs do not predict dimerization. Critically, although some LPXTG-anchored orthologs such as ScpC possess Ig-like domains, AlphaFold 3 predictions do not predict dimerization for these proteins either. This suggests that the specific arrangement of Ig domains found in the ScpH subfamily, rather than the mere presence of Ig folds, is required for the formation of the interlocked dimer observed here. The FIVAR domains, in contrast, do not contribute to dimerization but instead mediate oligomerization through interface III and interaction with eDNA, as discussed below. AlphaFold 3 predictions of ScpH homologs from multiple *Streptococcus* species that share Ig and FIVAR domains consistently yield high-confidence dimers (ipTM = 0.69-0.74) with a conserved butterfly-shaped arrangement similar to the experimentally determined dimer for ScpH (Fig. S7 and Fig. S8). Among these, two distinct subfamilies emerge. The first, exemplified by *S. sanguinis*, lacks an LPXTG motif entirely and instead terminates with FIVAR domains and an unfolded domain resembling the DNA-binding tail observed in ScpH; this group includes *S. gordonii*, *S. oralis*, *S. intermedius*, *S. salivarius*, *S. anginosus*, and *S. rubneri* (Fig. S7). The second subfamily retains a canonical LPXTG cell wall anchoring motif but also possesses FIVAR domains; this group includes *S. canis*, *S. equinus*, *S. gallolyticus*, *S. hillyeri*, *S. suis*, and *S. suivaginalis* (Fig. S8). Strikingly, members of this second group are also predicted to dimerize with comparable confidence, suggesting that the capacity for dimerization is a shared property of FIVAR-containing C5a peptidase homologs regardless of the presence of an LPXTG anchor.

Second, the capacity of ScpH dimers to oligomerize into an extended filament through the additive contributions of interfaces II, III, and IV is without precedent for this protease family. Interfaces II (Fn1-Fn1) and III (FIVAR2-FIVAR2) ensure longitudinal propagation along each row, while interface IV (protease-Ig1) provides lateral cohesion between the two rows. Although each of these interfaces is individually modest in buried surface area, their repetitive nature along the filament creates a cooperative stabilization mechanism in which the assembly grows increasingly stable with each added dimer. The Ig1 domain plays a particularly central role in this architecture, acting as a structural keystone by participating in both the intra-dimer interface I and the inter-row interface IV. AlphaFold 3 predictions of the FIVAR tails from homologs in the LPXTG-lacking subfamily (*S. gordonii*, *S. oralis*, *S. intermedius*, *S. salivarius*, *S. anginosus*, and *S. rubneri*) reveal that these domains dimerize with high confidence, indicating that interface III is conserved across this group and suggesting that these homologs are also capable of oligomerization (Fig. S9). By contrast, AlphaFold 3 does not predict FIVAR dimerization for the LPXTG-containing subfamily, consistent with the idea that these homologs function as cell wall-anchored dimers without the capacity for further oligomerization into extended filaments.

Third, and perhaps most remarkably, the ScpH filament is assembled on a scaffold of eDNA rather than being covalently anchored to the cell wall. The absence of an LPXTG motif, which is invariably present in the characterized orthologs ScpA (*S. pyogenes*), ScpB (*S. agalactiae*), and ScpC (*S. pyogenes*), clearly implied a fundamentally different mode of surface presentation. In the filament, this anchoring function is replaced by the FIVAR3 domain and the DNA-binding tail, which form an electropositive cleft that interacts with the negatively charged DNA backbone. EMSA experiments confirm that FIVAR3 together with the DNA-binding tail is required for DNA binding, while FIVAR3 alone is insufficient. Given the double-stranded nature of the ScpH filament, in which two rows of protomers flank a central DNA strand, repeated over potentially many DNA molecules, this architecture is likely to generate a mesh-like network of protease-decorated DNA filaments rather than an oriented linear assembly. Analysis of the electrostatic surface properties of the FIVAR and ND domains across the LPXTG-lacking subfamily reveals a conserved electropositive character similar to that of *S. sanguinis* ScpH, suggesting that these homologs also have the capacity to bind eDNA (Fig. S10). In the LPXTG-containing homologs, dimerization is confidently predicted by AlphaFold 3, but the absence of an electropositive DNA-binding surface suggests that these proteases do not interact with eDNA (Fig S11).

The observation that a putative C5a peptidase assembles directly onto eDNA places this system at the intersection of two active fields: biofilm matrix biology and immune evasion. eDNA is a critical structural component of streptococcal biofilms, providing mechanical stability and mediating intercellular cohesion^37^. Several of the species harboring ScpH homologs in the LPXTG-lacking subfamily, including *S. sanguinis*, *S. gordonii*, and *S. oralis*, are early colonizers of the oral cavity that actively release eDNA into the biofilm matrix^38–40^. In *S. sanguinis* and *S. gordonii*, eDNA release occurs through a specific, autolysis-independent mechanism driven by hydrogen peroxide produced by the pyruvate oxidase SpxB^39,40^, and biofilm formation by *S. sanguinis* is highly dependent on eDNA production^2,26^. The assembly of a protease onto this eDNA matrix represents a form of enzymatic functionalization of the biofilm, transforming a passive structural scaffold into an active immune evasion platform. Recent work has highlighted the concept of the "catalytic biofilm matrix"^34^, in which enzymatically active proteins are retained within the extracellular matrix through interactions with structural components such as exopolysaccharides, where they contribute to immune defense and nutrient acquisition^32^. The ScpH-eDNA filament described here provides a striking parallel in Gram-positive bacteria, with DNA rather than polysaccharide serving as the organizing scaffold.

Although the catalytic triad (Asp-His-Ser) characteristic of S8 subtilisin-like serine proteases is clearly resolved in the active site and the catalytic geometry is conserved upon superposition with *S. pyogenes* ScpA, we have not detected proteolytic activity *in vitro* under the conditions tested. The presence of a co-purified peptide in the active site suggests that ScpH has been in contact with a substrate or inhibitor during purification. Whether ScpH retains the ability to cleave complement components (C5a, C3, C3a) and pro-inflammatory cytokines (IFN-γ, IFN-λ) like its *S. pyogenes* ortholog, or has evolved to target different substrates, remains an open question. Notably, a recent study has shown that ScpH is highly abundant in *S. sanguinis* EMVs and that its deletion abolishes the ability of these vesicles to inhibit host interferon signaling, providing indirect evidence that ScpH is functionally active in an extracellular context^29^. Resolving the substrate specificity of ScpH will require systematic biochemical characterization with a panel of complement components, chemokines, and interferons relevant to the oral and cardiovascular niches.

The ScpH-eDNA filament adds a new dimension to the already multifaceted immune evasion strategy of *S. sanguinis*. This organism deploys H₂O₂ production to kill neutrophils, a cell wall-anchored nuclease to escape NETs, and functional amyloid fibrils to resist complement-mediated clearance. The eDNA-ScpH filament described here would constitute an additional, spatially distinct layer of defense, operating within the biofilm matrix itself. Remarkably, the H₂O₂ produced by SpxB serves a dual role in this context: it is both the bactericidal agent that kills competing species and neutrophils, and the trigger for the autolysis-independent release of eDNA that serves as the scaffold for ScpH assembly. This mechanistic coupling suggests a coordinated program in which a single metabolic output simultaneously generates a chemical weapon and the structural platform upon which an enzymatic defense is deployed. In the context of infective endocarditis, where *S. sanguinis* must survive in the bloodstream and colonize damaged cardiac endothelium under intense immune surveillance, such a multi-layered strategy could provide critical protection during the early stages of infection. The ScpH-eDNA filament may thus represent not only a novel biofilm functionalization mechanism but also a virulence architecture with direct clinical relevance.

## Material and Methods

### Strains and growth conditions

*S. sanguinis* strains were grown on plates containing Todd Hewitt (TH) broth (Difco) and 1% agar (Difco), or in liquid culture in THTH, i.e., TH broth containing 0.05% Tween 80 (Merck) to limit bacterial clumping, and 100 mM HEPES (Euromedex) to prevent acidification of the medium. Plates were incubated at 37 °C in anaerobic jars (Oxoid) under anaerobic conditions generated using Anaerogen sachets (Oxoid), while liquid cultures were grown statically in closed bottles within a standard atmospheric incubator (no added CO_2_), resulting in oxygen-limited conditions.

*Escherichia coli* (*E. coli*) strains 10β and BL21(DE3) (both from New England Biolabs) were used for cloning and recombinant protein expression, respectively. Specifically, the *E. coli* BL21(DE3)_SCPH_FIVAR3 and *E. coli* BL21(DE3)_SCPH_FIVAR3&ND strains were generated by transforming competent *E.coli* BL21(DE3) cells with the corresponding sequence-verified pET20b(+) expression plasmids. The construction and production of these plasmids are detailed in the following section. Bacteria were grown in liquid or on solid Luria-Bertani (LB) medium supplemented with ampicillin (100 µg/mL). Liquid cultures were incubated at 37 °C with orbital shaking at 180 rpm.

All *S. sanguinis* mutant strains were generated in the 2908 wild-type background^41^. The SS56 strain used in this study to purify ScpH filaments (in which the *pil* locus encoding T4P was entirely deleted) was previously described elsewhere^42^. The strain expressing the ScpH-ALFA-tag fusion was constructed using a previously described splicing PCR (sPCR) mutagenesis strategy^43^, starting from a constructed intermediate strain that expressed a ScpH-mChartreuse fusion. Briefly, this intermediate strain was generated by amplifying three overlapping fragments (upstream, mChartreuse insert, and downstream) using wild-type 2908 genomic DNA and a plasmid encoding the mChartreuse fluorophore^44^ as templates. These three amplicons were then fused via sPCR. Subsequently, the mChartreuse sequence was replaced by an ALFA-tag. Using the genomic DNA of the mChartreuse-expressing intermediate strain as a template, two overlapping fragments were amplified and assembled via sPCR. The ALFA-tag sequence was directly incorporated into the overlapping tails of the internal primers during amplification, effectively replacing the mChartreuse sequence. The bacterial strains and primers used in this study are detailed in Supplementary Tables S1 and S2, respectively.

### Plasmid construction for *E. coli* expression strains

The expression plasmids pET20b(+)_FIVAR3&ND and pET20b(+)_FIVAR3 were generated by PCR using Phusion High-Fidelity DNA Polymerase (New England Biolabs). Specifically, the pET20b(+)_FIVAR3&ND plasmid was first amplified from the parent vector prtS_antenna-pET20b(+), which contains the C-terminal sequence of ScpH (residues 1197-1506). Once sequence-verified, this pET20b(+)_FIVAR3&ND plasmid served as the template for a second PCR to construct the shorter pET20b(+)_FIVAR3 plasmid. All amplification products were digested with *DpnI* for 1 h at 37 °C to remove the template, purified using the NucleoSpin PCR Clean-up kit (Macherey-Nagel), and 5’-phosphorylated with T4 Polynucleotide Kinase overnight at 37 °C. Following an overnight ligation at room temperature with T4 DNA ligase, the plasmids were transformed into *E. coli* 10β competent cells via heat shock (42 °C, 30 s) and plated on LB agar supplemented with 100 μg/mL ampicillin.

Single colonies were selected and grown overnight at 37 °C with shaking (180 rpm) in liquid LB-ampicillin. Plasmids were isolated using the NucleoSpin Plasmid EasyPure kit (Macherey-Nagel) and verified by sequencing (Eurofins). For recombinant protein production, the sequence-verified plasmids were subsequently transformed into the expression strain *E. coli* BL21(DE3) using the same heat-shock procedure and selected on LB-ampicillin agar plates overnight at 37 °C. All plasmids used in this study are listed in Supplementary Table S3.

### Expression and purification of recombinant ScpH proteins

Recombinant ScpH_FIVAR3&ND and ScpH_FIVAR3 proteins, both harboring an N-terminal Strep-Tag II, were expressed in *E. coli* BL21(DE3). Cultures (2 L) were grown to an OD600 of 0.5, induced with 1 mM IPTG, and incubated overnight at 16 °C. Cells were harvested by centrifugation (8,000 × g, 30 min) and resuspended in lysis buffer (20 mM Tris-HCl, 50 mM NaCl, pH 7.4) supplemented with 10 mg lysozyme, DNase, and a cOmplete^TM^ protease inhibitor cocktail tablet (Roche). Cells were mechanically disrupted by high-pressure homogenization (EmulsiFlex, Avestin), and the lysate was clarified by centrifugation to recover the soluble protein fraction.

Protein purification was performed by affinity chromatography using an ÄKTA Avant automated system equipped with a 5 mL StrepTactin-XT column (Cytiva). The clarified lysate was loaded onto the pre-equilibrated column, followed by a stringent wash with 10 column volumes (CV) of high-salt wash buffer (20 mM Tris-HCl, 3 M NaCl, pH 7.4) to remove non-specific contaminants. The column was subsequently washed with 10 CV of equilibration buffer (20 mM Tris-HCl, 50 mM NaCl, pH 7.4) to return to low-salt conditions prior to elution. Bound proteins were eluted using commercial Buffer XT (IBA) containing desthiobiotin. The purity of the eluted fractions was assessed by 10% SDS-PAGE (samples were denatured at 95 °C for 5 min prior to loading) and visualized by Coomassie blue staining (Fig S1b and S1c).

### Electrophoretic mobility shift assay (EMSA)

Protein-DNA interactions were assessed *in vitro* by co-incubating 200 ng of a 1297 bp DNA fragment with 20 µg of recombinant FIVAR3 & ND or FIVAR3 protein for 1 h at 37 °C. The resulting protein-DNA complexes were resolved by electrophoresis on a 1% (w/v) agarose gel in TAE buffer supplemented with SYBR^TM^ Safe DNA gel stain (Invitrogen). Electrophoresis was performed at 110 V for 50 min, and the gels were visualized under UV illumination using a ChemiDoc imaging system (Bio-Rad).

### ScpH-DNA filament purification

*S. sanguinis* SS56 pre-cultures were inoculated at an initial OD600 of 0.02 into THTH, and grown statically at 37 °C to an OD600 of 0.35-0.40 (4 × 1 L). Cells were initially harvested by centrifugation (4,500 x g, 4 °C), and the loose pellets were carefully pooled using wide-bore pipettes, followed by a second centrifugation step (7,000 x g, 10 min, 4 °C). After the residual supernatant was completely discarded, each cell pellet (equivalent to 1 L of culture) was resuspended in 1.5 mL of cold THTH via repeated, controlled pipetting until the initially viscous suspension became fluid. Intact cells and large debris were removed by two sequential clearing centrifugations (10,000 x g, 10 min, 4 °C), and the resulting clarified supernatant was subjected to ultracentrifugation (100,000 × *g*, 1 h, 4 °C) to pellet the ScpH filaments. Following supernatant removal and a brief spin, to collect the residual buffer (∼40 µL total), this minimal volume was used to serially and gently resuspend the pooled filament pellets. The resulting highly concentrated filament suspension was subsequently used for the preparation of cryo-electron microscopy (cryo-EM) grids, while the remainder was analyzed by SDS-PAGE. Protein bands were subsequently excised and identified via mass spectrometry as described in Fig. S1a.

### Cryo-EM sample preparation and data acquisition

R2/2 Cu 200 mesh grids (Quantifoil) were glow-discharged for 40 s at 2.7 mA. Then, 5 µl of freshly purified filaments were applied and the excess of sample was immediately blotted away (3.5 - 4s blot time, 4 °C chamber temperature, 100% humidity) in a Vitrobot Mark IV (Thermo Fisher Scientific), before being plunge-frozen in liquid ethane. Cryo images of the purified filaments were recorded with a Glacios 2 microscope (Thermo Fisher Scientific) operated at 200 kV and equipped with a Falcon 4i camera and a SelectrisX energy filter (Thermo Fisher Scientific). Dose fractioned data were collected in a defocus range of −0.2 to −2.4 µm at 130,000x magnification, corresponding to a pixel size of 0.885 Å, using Smart EPU software (Thermo Fisher Scientific). The total dose of electrons was 51.80 e^-^/Å^2^, with 1.72 e^-^/Å^2^ per frame. The cryo-EM data collection statistics are detailed in Supplementary Table S4.

### Image processing

All data processing was performed in cryoSPARC^45^, with the overall workflow outlined in Fig. S3. Briefly, 25,079 movie stacks underwent motion correction and CTF estimation, yielding 22,037 usable micrographs. Particle picking was executed using Topaz^46^ via an iterative training loop: an initial model based on manual picking was continuously retrained using the best 2D classes from subsequent extractions. These extractions from the micrographs were performed using a box size of 512px. Following 2D classification, this process yielded 202,815 particles, which were subjected to *ab initio* reconstruction and heterogeneous refinement. A cleaned subset of 77,124 particles underwent a second identical round to further remove heterogeneity. The resulting 68,107 particles were processed via C1 non-uniform (NU) refinement, local CTF refinement, and reference-based motion correction. A subsequent C1 NU-refinement generated a 3D volume from 65,221 particles. From this volume, three parallel refinement strategies were applied: (i) a direct C1 local refinement focused on the DNA region, (ii) a re-extraction of particles from the micrographs using a larger box size of 900 px, followed by *ab initio* reconstruction (yielding 52,657 particles) and a final C1 homogeneous refinement, and (iii) a C2 NU-refinement, which yielded the consensus map of the complex. This consensus map was subsequently subjected to C2 symmetry expansion, producing 130,442 pseudo-particles for a final targeted local refinement. This high-resolution C2-expanded local map was subsequently used for atomic model building.

### Protein structure prediction and sequence sourcing

Structural prediction of the *Streptococcus sanguinis* ScpH protein (strain 2908, ENA accession PRJEB7884)^41^ and its streptococcal homologs were performed using the AlphaFold 3 server^47^. All sequences and their corresponding accession numbers used in this study are detailed in Supplementary Table S5.

For *S. sanguinis* ScpH, the modeled sequence excluded the N-terminal pre/pro domain, thus starting directly at residue 176. The predicted overall fold of the ScpH homodimer, as well as a single monomeric chain extracted from this dimeric model, are presented in Fig. S5. To investigate specific regions of ScpH, the flexible C-terminal tail (residues 1281–1506, comprising FIVAR 1–3 domain and the DNA-binding tail was modeled as a dimer both in isolation and in complex with a 45-bp double-stranded DNA molecule (5’-ACCGGGCCACTCAAAAGTCATTTTGTAATTGTACACGATGTGATG-3’) are presented in Fig S5. Furthermore, ScpH homodimers were modelled in the presence of human C5a (residues 678–751, UniProtKB P01031). The predicted folds for the ScpH homodimer with C5a and the extracted C5a monomer are presented in Fig. S5.

For structural comparison, homologs were identified by an NCBI BLASTp^48^ search against the *Streptococcus* genus using the *S. sanguinis* ScpH C-terminal region (residues 1105–1506). Full-length S8 family serine peptidases were selected and evaluated for the LPXTG motif using InterProScan^49^. Fourteen representative sequences were subsequently modeled with AlphaFold 3^47^: seven lacking the LPXTG motif (*S. anginosus*, *S. gordonii*, *S. intermedius*, *S. oralis*, *S. rubneri*, *S. salivarius*) and seven containing it (*S. canis*, *S. equinus*, *S. gallolyticus*, *S. hillyeri*, *S. suis*, *S. suivaginalis*). Consistent with the ScpH modeling strategy, the N-terminal pre/pro-domains of all selected homologs were excluded prior to structure prediction. Additionally, for the homologs lacking the LPXTG motif, the isolated C-terminal tail (comprising the FIVAR 1–3 domains) was also modeled as a dimer to allow direct structural comparison with the ScpH tail.

### Atomic model building

The full dimeric models of ScpH generated by AlphaFold3 were initially rigid-body docked into a cryo-EM map sharpened with LocScale 2^50^ using UCSF ChimeraX^51^. Subsequently, a map sharpened by EMReady2^52^ was used to accurately rebuild and fit the protein backbone. Regions of the AlphaFold models lacking corresponding map density were manually deleted. The trimmed model was then subjected to iterative rounds of refinement against a map sharpened in PHENIX (*phenix.autosharpen*)^53^. This optimization process alternated between manual adjustments in ISOLDE^54^ and automated real-space refinement using *phenix.real_space_refine*^55^. Finally, the atomic model was validated using *phenix.validation_cryoem*^56^ and the wwPDB validation server^57^. To provide a comprehensive structural representation of the mature ScpH, a composite model was generated by grafting missing flexible domains from the initial AlphaFold 3 prediction onto the refined atomic model. Detailed statistics for the cryo-EM data refinement and the model-to-map Fourier shell correlation (FSC) plot are provided in Supplementary Table S4 and Fig. S12, respectively.

### Interface evaluation and comparative analysis

Macromolecular interfaces and interaction surfaces within the ScpH dimeric complexes (Fig. 3) were evaluated using the PDBe PISA (Proteins, Interfaces, Structures and Assemblies) server. Specifically, PISA was utilized to quantify the buried surface areas (Å²), and identify the specific non-covalent interactions (hydrogen bonds, salt bridges, and hydrophobic contact) stabilizing the assembly across the distinct domain interfaces (denoted I to IV).

Structural comparison analyses and RMSD (Root Mean Square Deviation) calculations were performed using UCSF ChimeraX. The ScpH atomic model was compared to *S. pyogenes* ScpA (PDB 3EIF). A global alignment was first executed using the *Matchmaker* tool, restricting the superposition to the conserved N-terminal core of ScpH (residues 176–1105) and applying an iterative pruning algorithm to exclude highly flexible regions from the final calculation. The geometric conservation of the active site was further evaluated by a local structural alignment of the alpha carbons (Cα) of the catalytic triad residues (Asp203, His269, Ser597 for ScpH; Asp130, His193, Ser512 for ScpA).

### Immunofluorescence

Bacterial biofilms were grown in 8-well chamber slides (μ-Slide 8 well, Ibidi). Wells were pre-coated with 0.01% poly-L-lysine overnight at 4 °C. Following a PBS wash, bacterial cultures were inoculated at an optical density (OD) of 0.1 and incubated under static conditions at 37 °C for 8 hours. The culture medium was then aspirated, and the biofilms were gently washed with 1X PBS.

Biofilm cells were fixed with 4% paraformaldehyde (PFA) for 1 hour at 37 °C, followed by quenching with 150 mM glycine in PBS for 1 hour at 37 °C. After three washes with 1X PBS, samples were blocked in a buffer containing PBS, 0.1% Tween-20, and 2% BSA for 15 minutes at 37 °C with gentle agitation (60 rpm). Biofilms were then co-incubated with primary antibodies for 1 hour at 37 °C (60 rpm): an Atto 488-conjugated anti-ALFA-tag alpaca nanobody (FluoTag-X2 anti-ALFA, NanoTag Biotechnologies; 1:500 dilution) and a mouse anti-DNA antibody (Abcam; 1:1000 dilution).

Following three washes with PBS supplemented with 0.1% Tween-20, samples were incubated with an Alexa Fluor 568-conjugated goat anti-mouse IgG secondary antibody (1:1000 dilution) for 1 hour at 37 °C (60 rpm) to detect the anti-DNA primary antibody. Unbound secondary antibodies were removed by three final consecutive washes with PBS prior to imaging.

Image acquisition was performed using a Leica DMI8 inverted microscope equipped with a 100× oil-immersion objective and an sCMOS camera. Fluorescence imaging was conducted using specific filter sets tailored for GFP, Atto 488, and Alexa Fluor 568. The instrument setup and image capture were controlled via the LAS X software (Leica Microsystems). Exposure times were optimized independently for each channel to maximize the signal-to-noise ratio while preventing pixel saturation.

## Data Availability

The cryo-EM maps have been deposited in the Electron Microscopy Data Bank (EMDB) under accession codes EMD-58287 (consensus map), EMD-58274 (local refinement, deck region, sharpened map used for atomic model building), EMD-58288 (local refinement, deck region, raw map), EMD-58289 (local refinement, DNA) and EMD-58299 (large box reconstruction). Atomic coordinates have been deposited in the Protein Data Bank (PDB) under accession code 31CH. Accession numbers for the sequences used in AlphaFold 3 predictions are provided in the corresponding figures. All other data supporting the findings of this study are available from the corresponding author upon reasonable request.

## Acknowledgements

We thank Axel Siroy for expert assistance at the cryo-EM platform of the IECB. This work was supported by the Agence Nationale de la Recherche (ANR ComPil, ANR-21-CE11-0008) and the Centre National de la Recherche Scientifique (CNRS).

## Author Contributions

R.A. performed cryo-EM data collection, image processing, model building, AlphaFold 3 predictions, and contributed to manuscript preparation including figure design. L.B. performed DNA binding experiments and A.V. performed fluorescence microscopy experiments, both under R.A.’s supervision. L.P. generated *S. sanguinis* strains. V.P. co-supervised the project. R.F. conceived and supervised the project, performed structural analysis, and wrote the manuscript with input from all authors.

---

**Figure S1:**
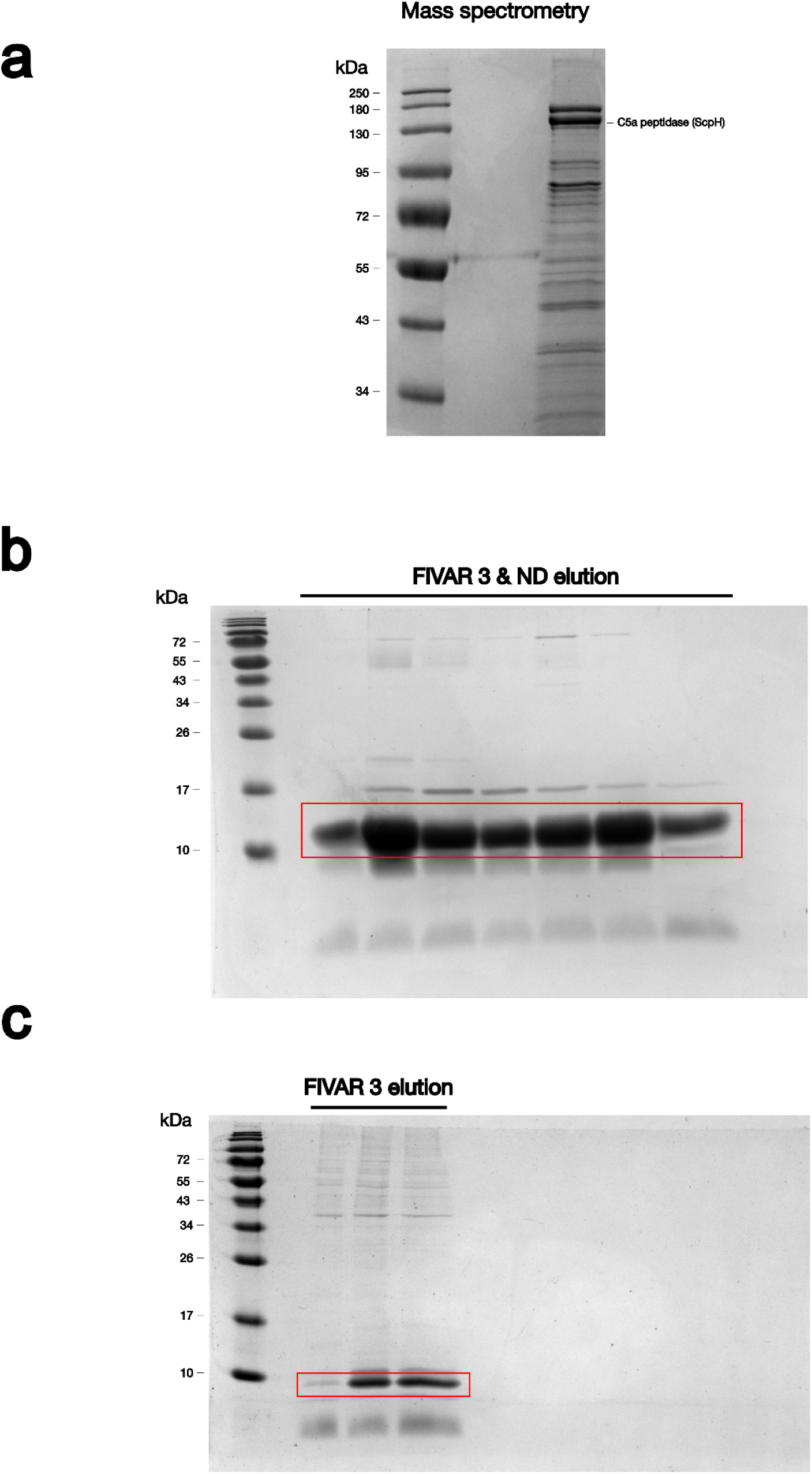
Identification of native ScpH and purification of recombinant C-terminal domains. **(a)** SDS-PAGE and mass spectrometry analysis of the sheared surface-associated material from *S. sanguinis*. The major high-molecular-weight protein band was excised and identified as a C5a peptidase homolog (ScpH). **(b, c)** Purity assessment of the recombinant ScpH C-terminal constructs used for *in vitro* DNA binding assays. SDS-PAGE gels visualized by Coomassie blue staining show the eluted fractions of the Strep-tagged FIVAR 3 & ND construct **(b)** and the FIVAR 3 domain alone **(c)** following affinity chromatography. The target protein bands corresponding to the purified recombinant proteins are highlighted by red boxes.

**Figure S2:**
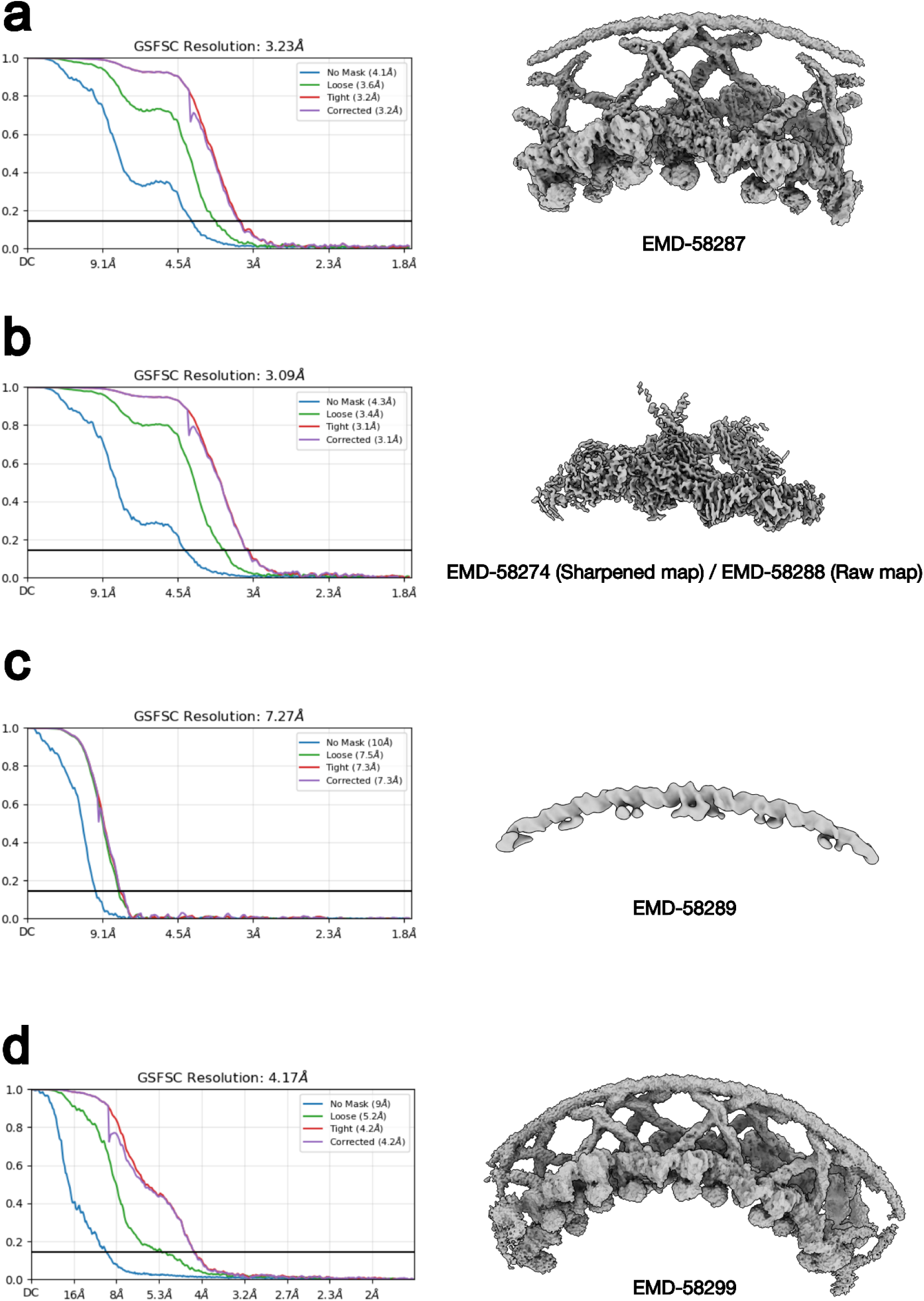
Cryo-EM density maps and Gold-Standard Fourier Shell Correlation (FSC) curves. Each panel displays the 3D reconstruction alongside its corresponding Gold-Standard FSC plot, with the reported global resolution determined at the 0.143 criterion (horizontal black line) of the corrected curve. **(a)** Consensus reconstruction of the ScpH assembly with C2 symmetry applied, yielding a 3.23 Å map. **(b)** Locally refined map focused on the structural deck region following C2 symmetry expansion, resolving to 3.09 Å. **(c)** Focused local refinement of the central DNA filament, reaching 7.27 Å resolution. **(d)** Global reconstruction of the extended filament, processed with a larger box size to encompass up to five consecutive dimers, resulting in a 4.17 Å map.

**Figure S3:**
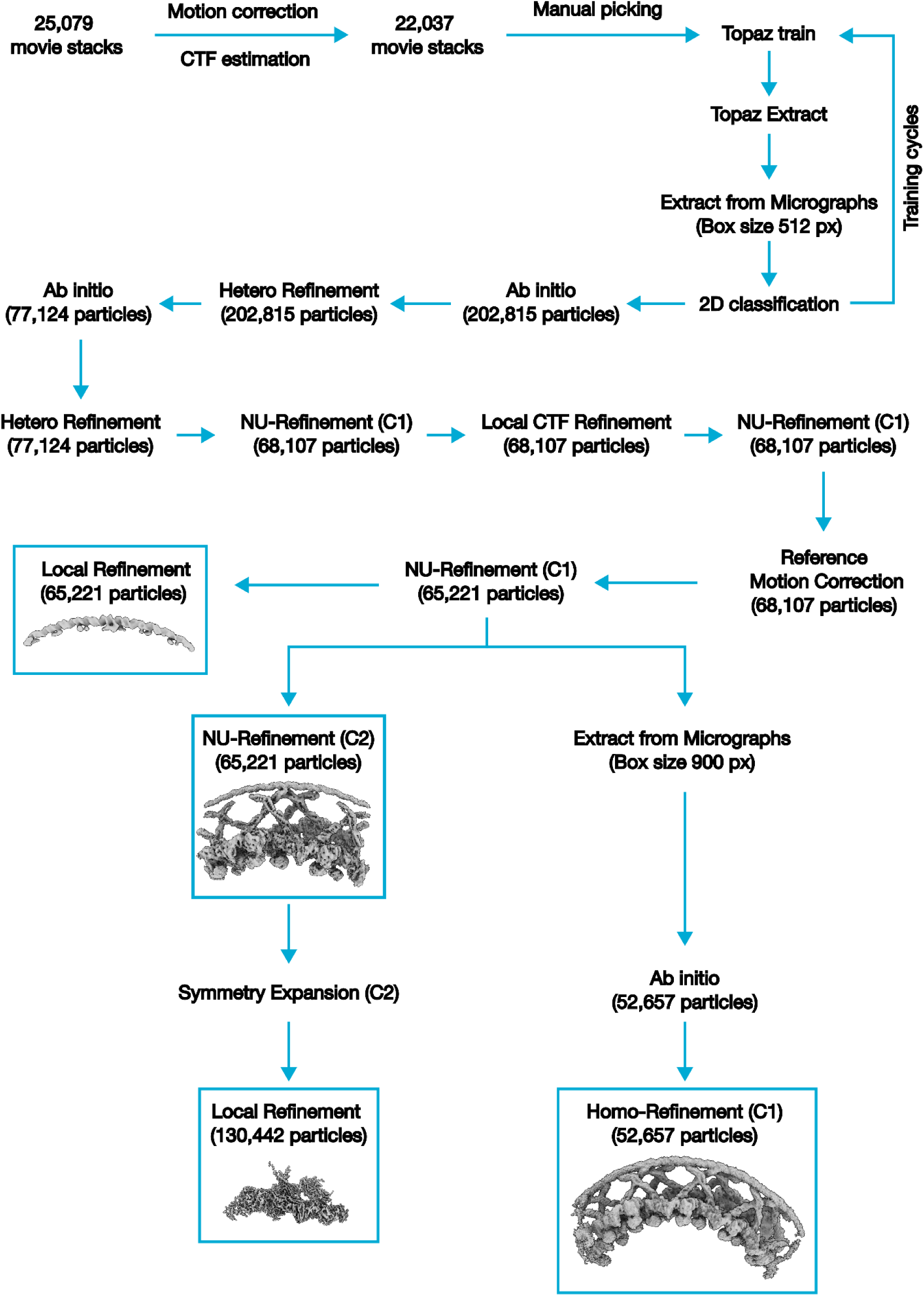
Cryo-EM data processing workflow. Flowchart illustrating the image processing and 3D reconstruction pipeline performed in cryoSPARC. The schematic summarizes the progression from raw micrograph processing and particle picking to the generation of the consensus map. It also outlines the three parallel refinement strategies (DNA-focused local refinement, C2 symmetry expansion for the deck region, and large-box particle re-extraction) used to obtain the final detailed and global reconstructions.

**Figure S4:**
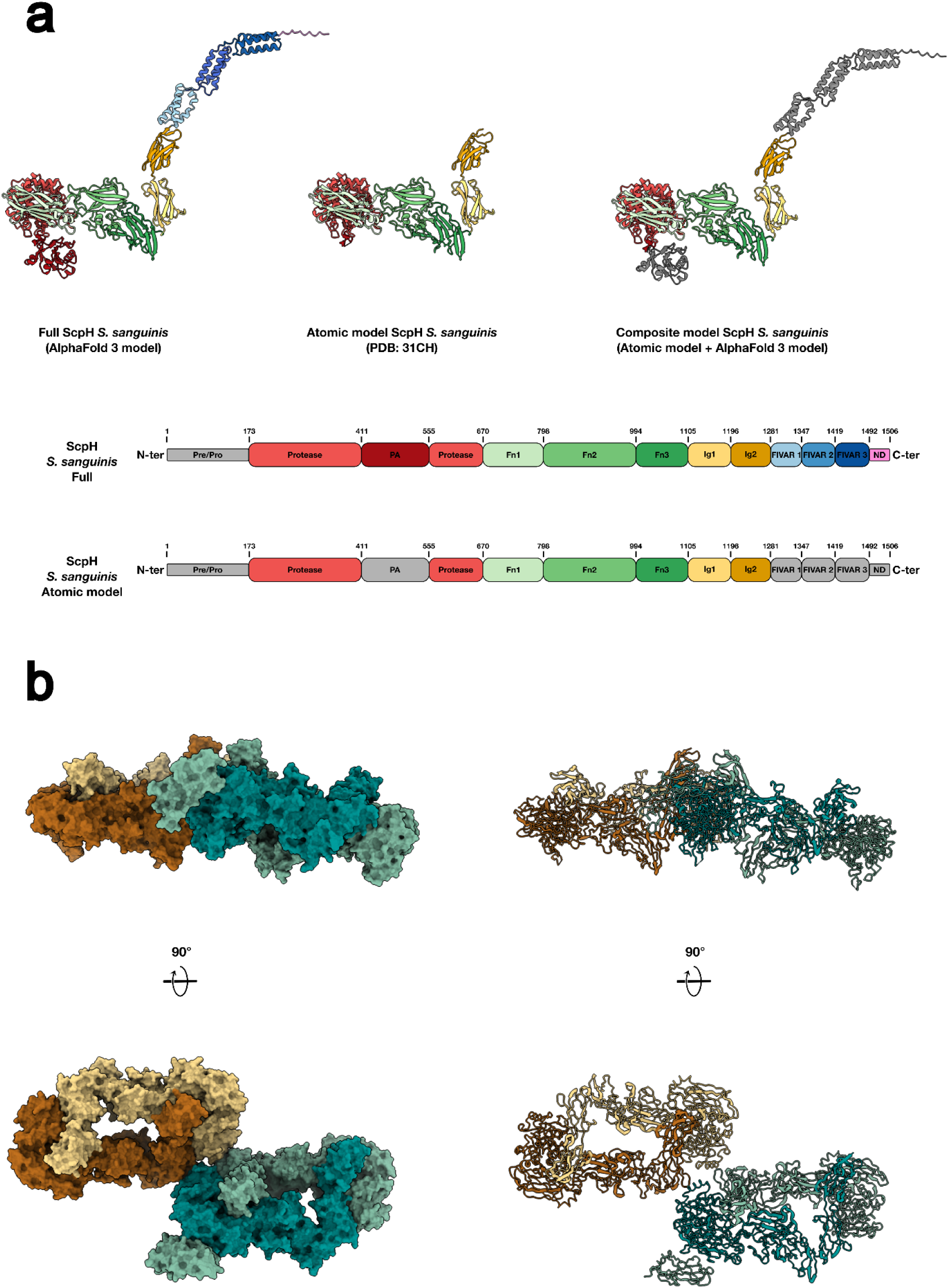
Atomic model building and domain organization of ScpH. **(a)** Domain boundaries and structural models of the ScpH monomer. The linear diagrams illustrate the domain organization of the full-length protein alongside the specific regions captured in the experimentally determined atomic model. Ribbon representations display the full AlphaFold 3 prediction, the atomic model built from the cryo-EM map, and the final composite model generated by grafting the predicted PA subdomain and flexible C-terminal arch (FIVAR and ND domains) onto the experimental structure. The linear diagrams and ribbon domains are colored according to the color scheme described in Figure 2. Note that regions shown in grey on the linear diagrams correspond to flexible domains that could not be modeled in the experimental atomic structure, and are therefore absent from the corresponding ribbon representations **(b)** Orthogonal views of the refined atomic model fitted into the locally refined C2-expanded cryo-EM map of the filament deck region, validating the experimentally determined structural framework for the inter-protomer contacts.

**Figure S5:**
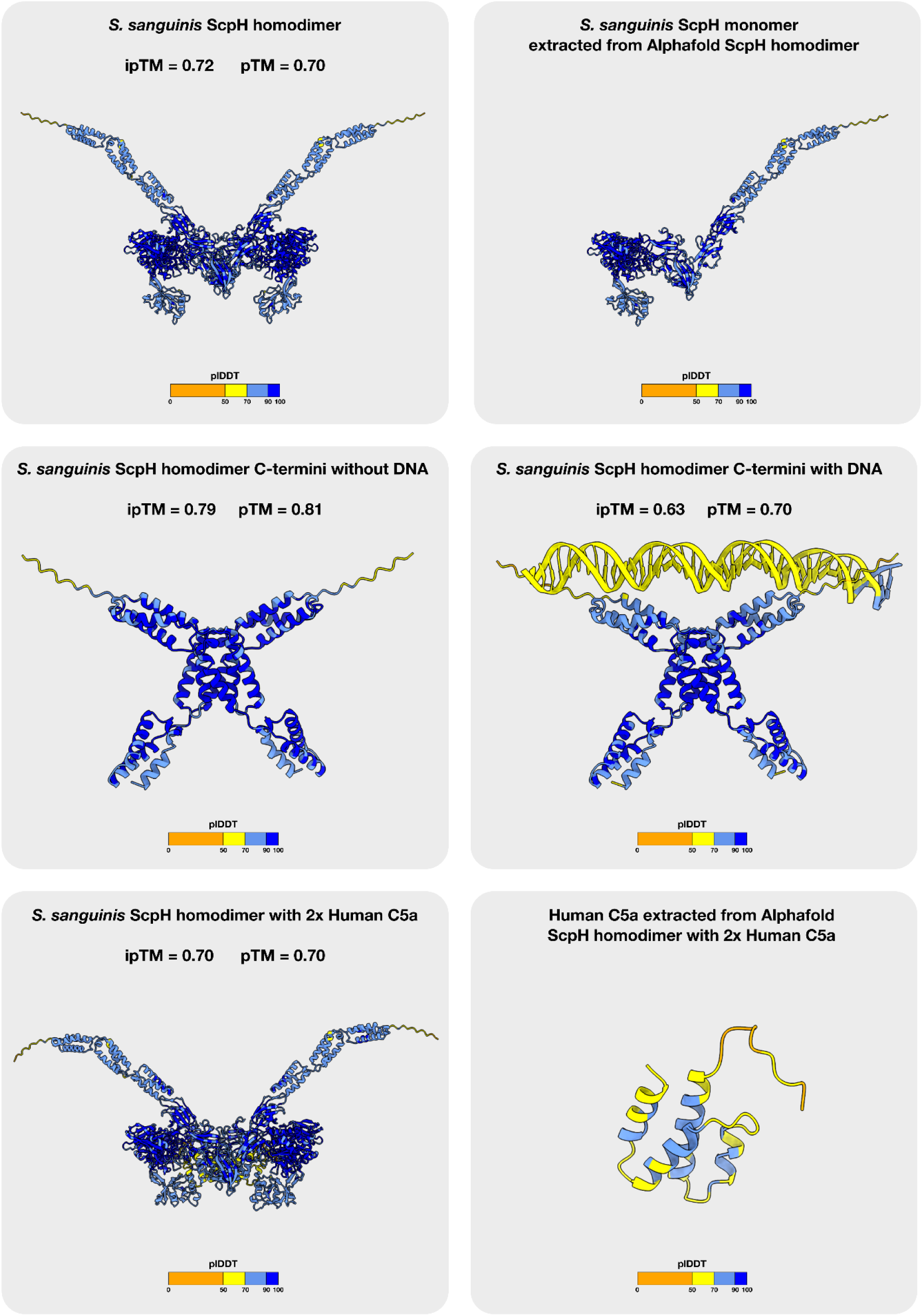
AlphaFold 3 structural predictions of ScpH and molecular complexes. Ribbon representations of the generated models are colored according to their predicted local distance difference test (pLDDT) confidence scores, ranging from very low (orange, <50) to very high (dark blue, >90). The panels display the prediction of the full-length ScpH homodimer alongside an extracted monomer, illustrating the global architecture. To investigate specific structural interactions, the flexible C-terminal region (comprising the FIVAR 1-3 and ND domains) was modeled both as an isolated dimer and co-folded with a 45-bp double-stranded DNA molecule, highlighting the formation of the DNA-receptive cradle. Additionally, the ScpH homodimer was co-folded with human C5a (shown with the full complex and as an extracted peptide) to assess the compatibility of the substrate-binding groove. Overall interface and structural confidence metrics (ipTM and pTM) are indicated for each corresponding prediction.

**Figure S6:**
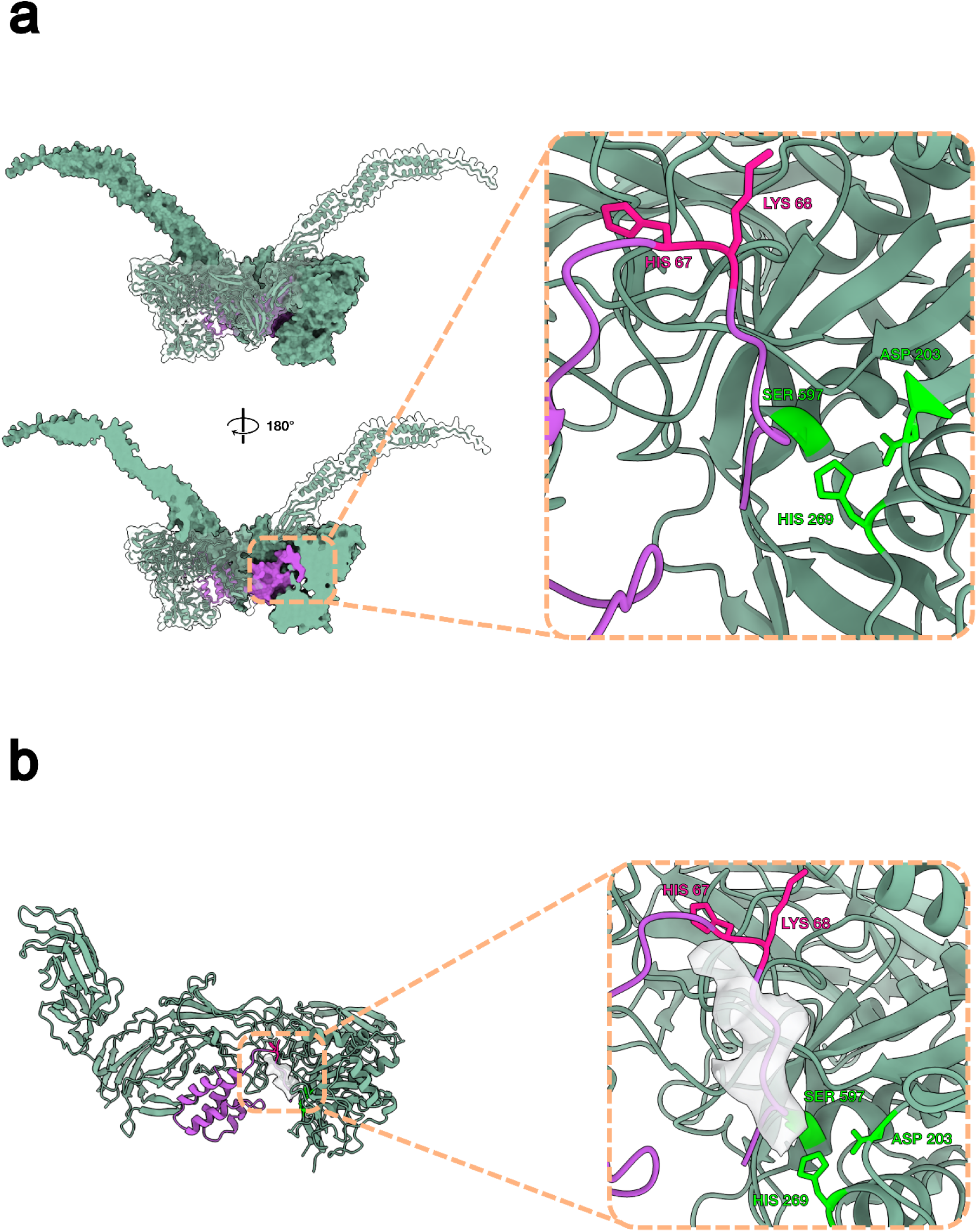
Structural compatibility of the ScpH active site with human C5a. **(a)** AlphaFold 3 co-fold prediction of the ScpH dimer in complex with human C5a. The predicted model positions the C5a peptide (purple) within the substrate-binding groove of the protease domain. A zoomed-in view reveals that the known C5a cleavage site (between His67 and Lys68, highlighted in pink) is located in close proximity to the ScpH catalytic triad (green). **(b)** Superposition of the AlphaFold 3-predicted ScpH-C5a complex with the experimental cryo-EM map. The structural alignment demonstrates that the predicted trajectory of the C5a cleavage site passes directly through the unmodeled density (white surface) observed in the active site cleft, illustrating that this experimental density occupies the expected substrate-binding path.

**Figure S7:**
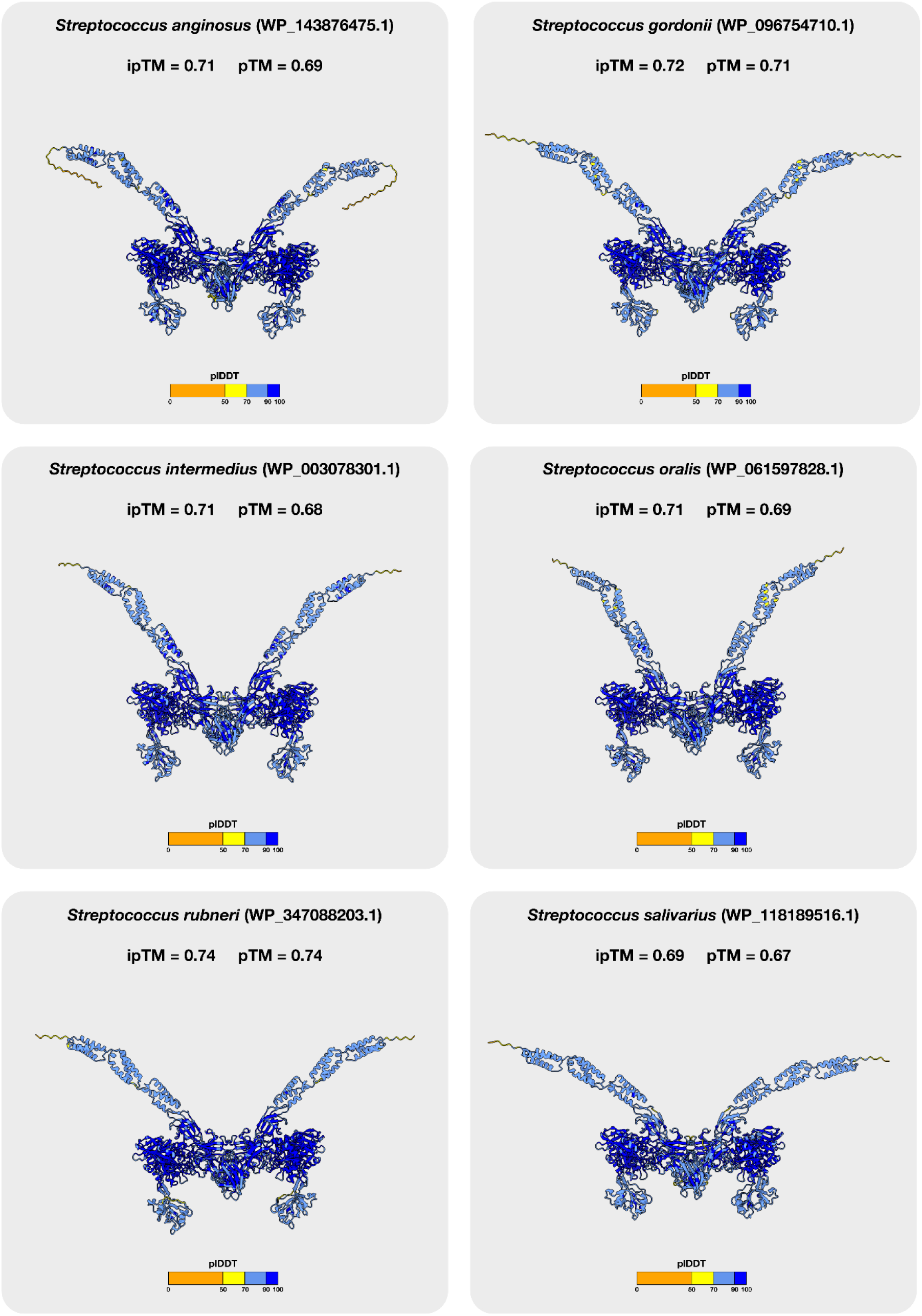
AlphaFold 3 structural predictions of ScpH homologs lacking the LPXTG motif. Ribbon representation of the predicted dimeric structures of various streptococcal ScpH homologs that naturally lack the canonical cell wall-anchoring sequence. The models are colored according to their pLDDT confidence scores. Notably, all these predictions consistently yield high-confidence dimers with a conserved butterfly-shaped architecture, closely mirroring the experimentally determined *S. sanguinis* ScpH dimer. Global interface and structural confidence metrics (ipTM and pTM) are indicated above each corresponding model.

**Figure S8:**
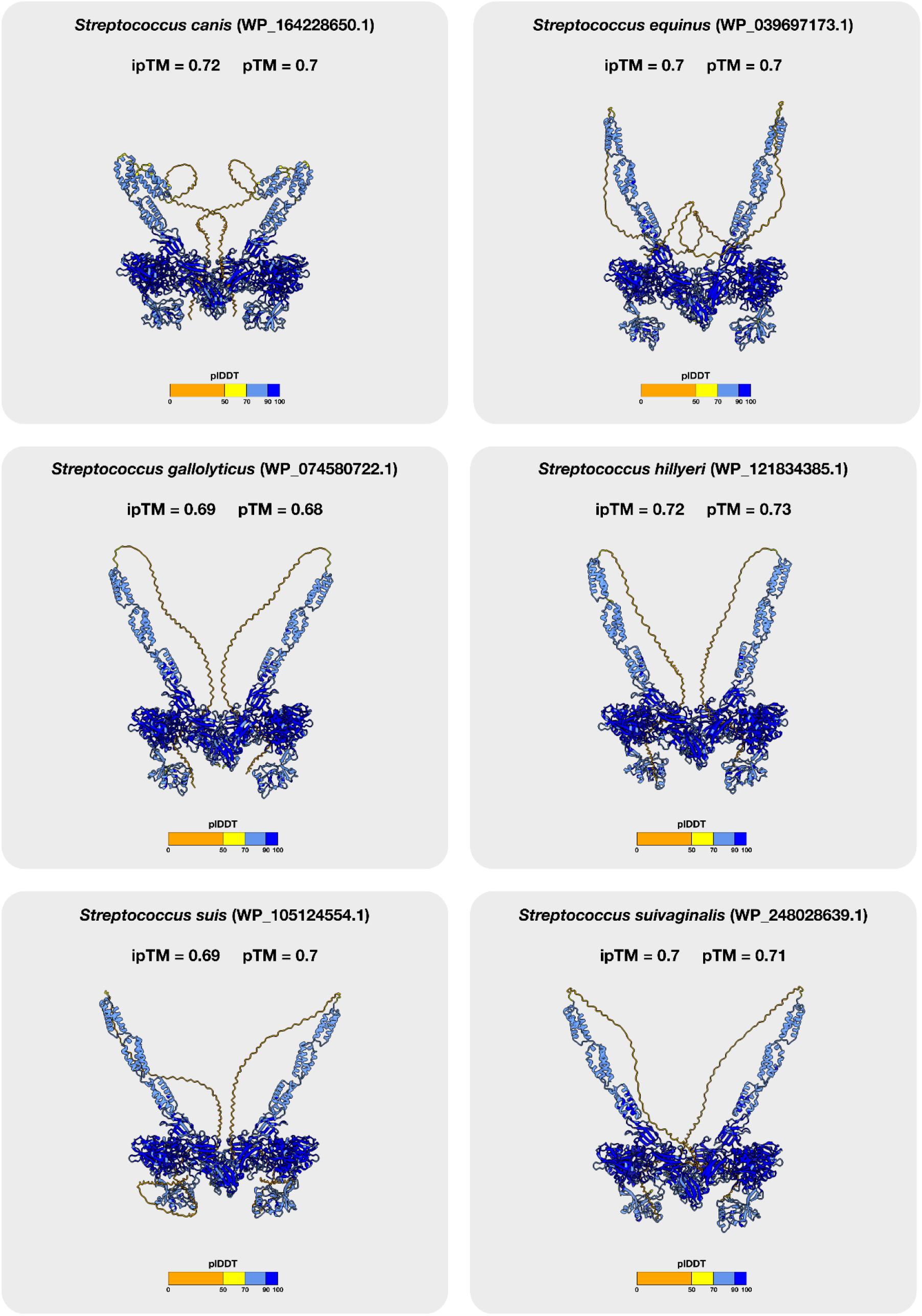
AlphaFold 3 structural predictions of ScpH homologs containing the LPXTG motif. Ribbon representation of the predicted dimeric structures of various streptococcal ScpH homologs that retain the canonical cell wall-anchoring LPXTG sequence. The models are colored according to pLDDT confidence scores. Notably, these predictions consistently yield high-confidence dimers, suggesting that the capacity for dimerization is a shared property of FIVAR-containing ScpH homologs regardless of the presence of an LPXTG anchor. Global interface and structural confidence metrics (ipTM and pTM) are indicated above each corresponding model.

**Figure S9:**
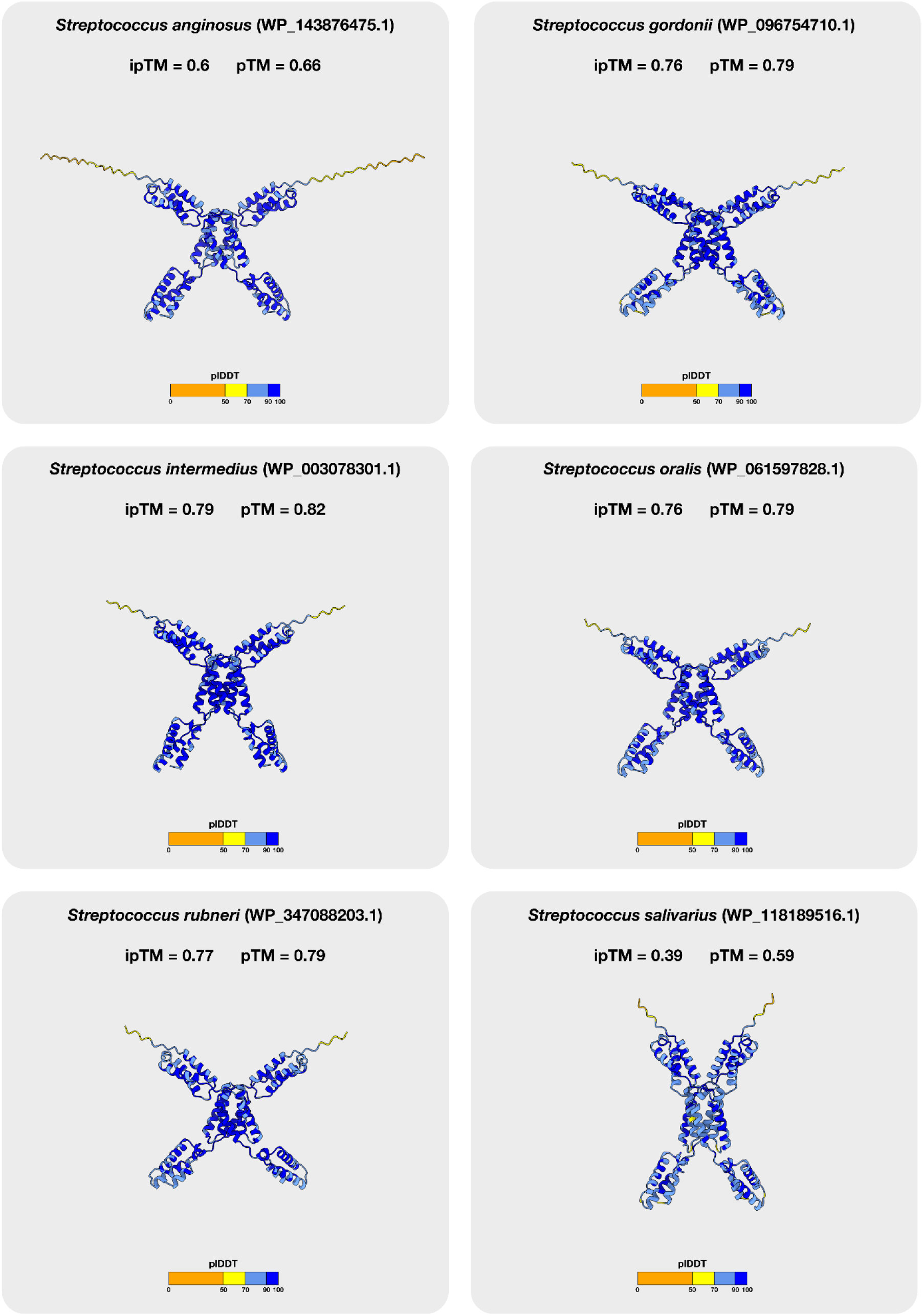
AlphaFold 3 structural predictions of isolated FIVAR tails from LPXTG-lacking ScpH homologs. Ribbon representation of the predicted dimeric structures of the isolated C-terminal tails (comprising the FIVAR domains) for ScpH homologs lacking the LPXTG motif. The models are colored according to their pLDDT confidence scores. These predictions consistently yield high-confidence dimers, indicating that the FIVAR-mediated dimerization interface (Interface III) is structurally conserved across this subfamily. This strongly suggests that these LPXTG-lacking homologs share the capacity for higher-order oligomerization into extended filaments. Global interface and structural confidence metrics (ipTM and pTM) are indicated above each corresponding model.

**Figure S10:**
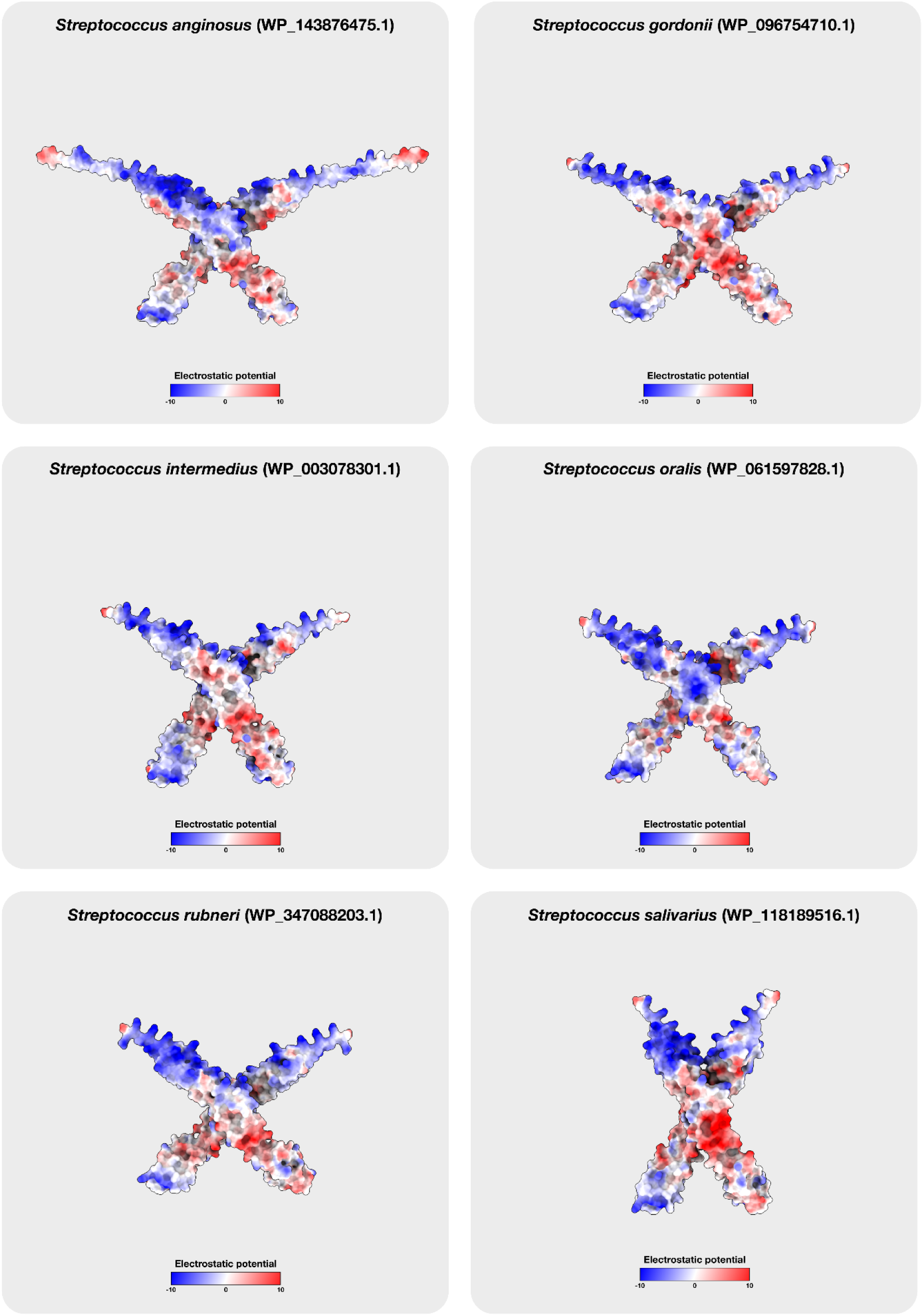
Surface electrostatic potential of the C-terminal tails from LPXTG-lacking ScpH homologs. Surface representations display the AlphaFold 3-predicted dimeric structures for the isolated C-terminal tails of ScpH homologs lacking the LPXTG motif. The models are colored according to their calculated electrostatic potential, ranging from negatively charged (red) to positively charged (blue). Similar to the *S. sanguinis* ScpH protein, this analysis reveals a strongly conserved electropositive character across this subfamily. This positively charged surface suggests that all these homologs share the capacity to bind eDNA through charge complementarity.

**Figure S11:**
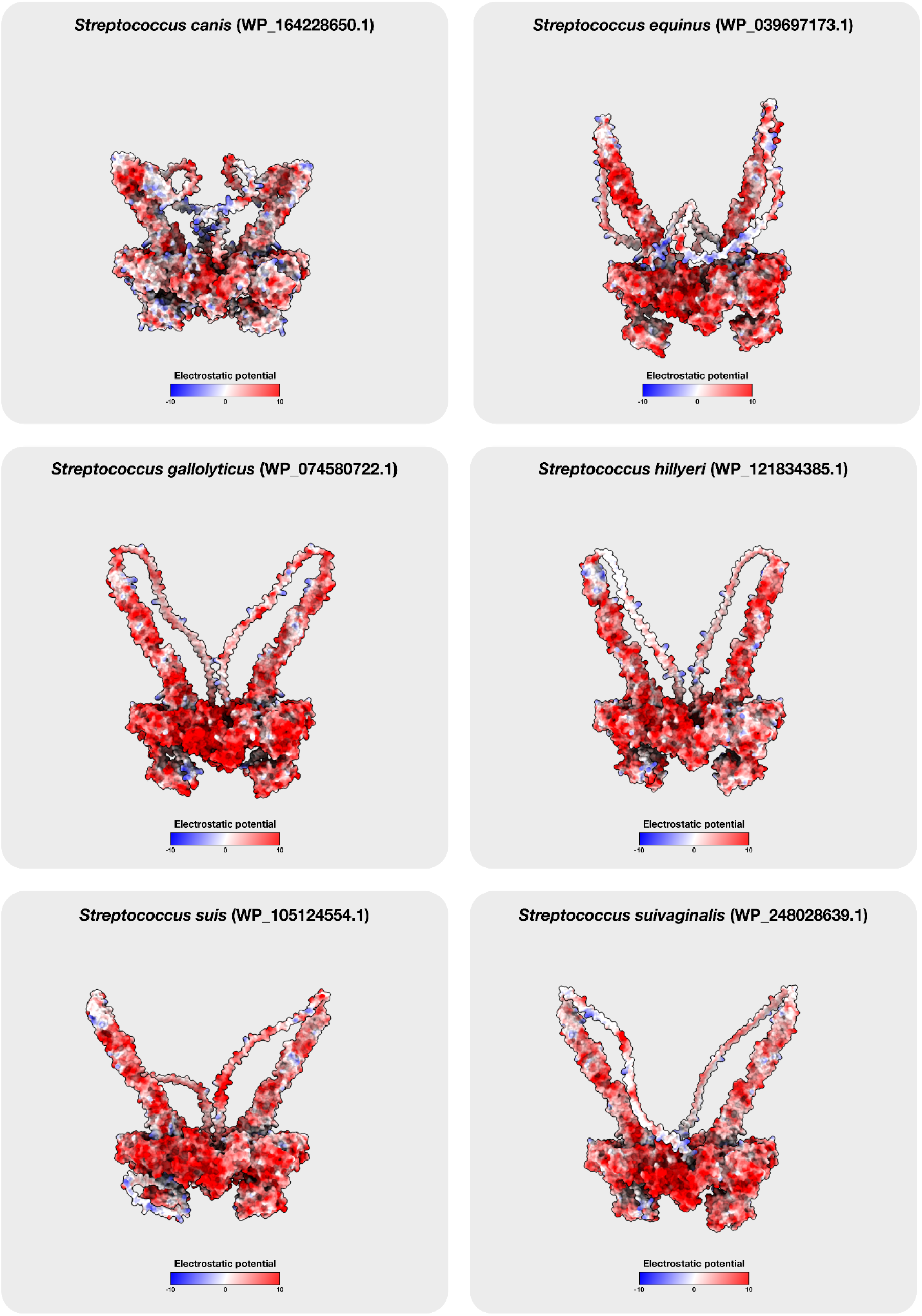
Surface electrostatic potential of ScpH homologs containing the LPXTG motif. Surface representations display the AlphaFold 3-predicted dimeric structures for streptococcal ScpH homologs that retain the canonical cell wall-anchoring LPXTG sequence. The models are colored according to their calculated electrostatic potential, ranging from negatively charged (red) to positively charged (blue). In contrast to the LPXTG-lacking homologs, these predictions reveal the absence of a strongly electropositive DNA-binding surface. This lack of charge complementarity suggests that these LPXTG-containing proteases do not interact with eDNA.

**Figure S12:**
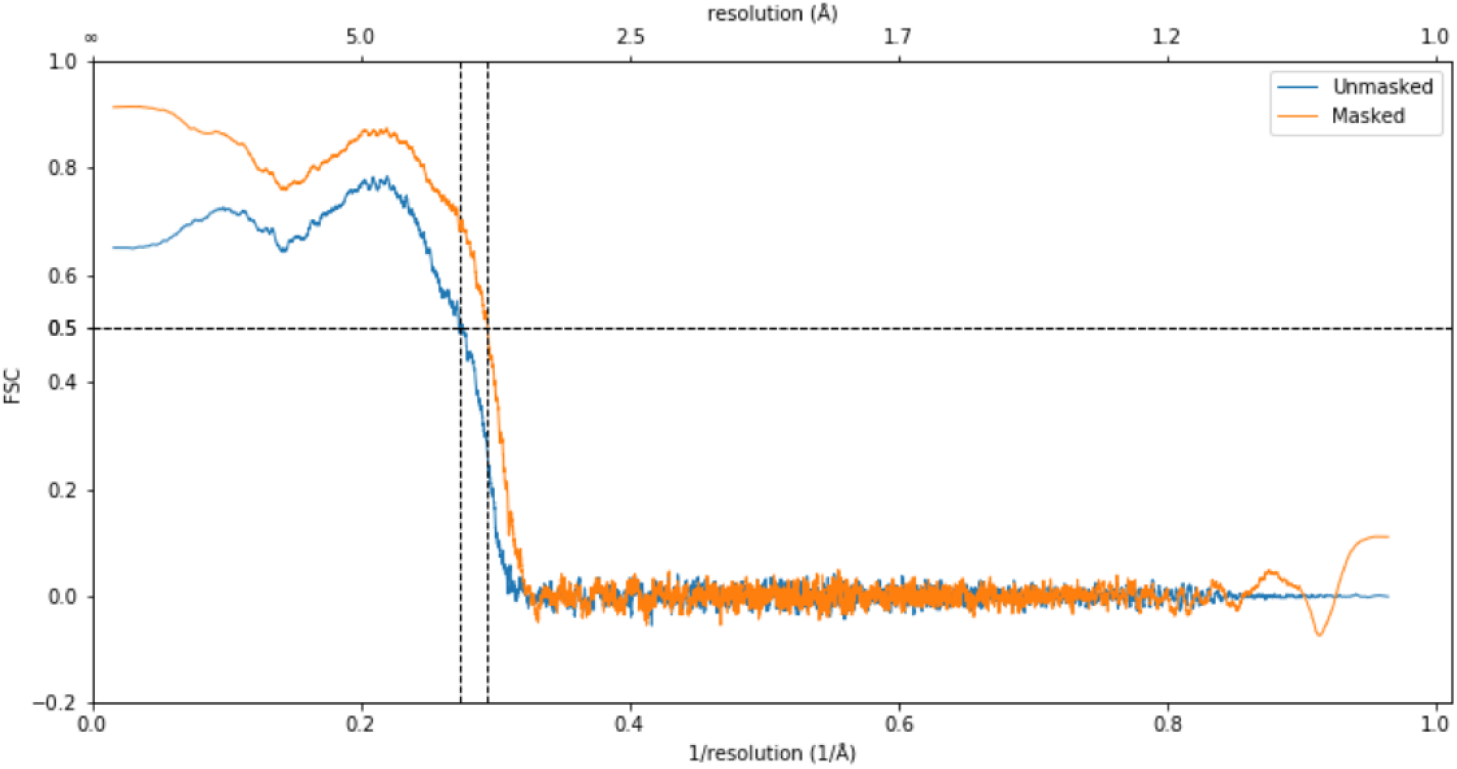
Model-to-map Fourier Shell Correlation (FSC) curve. The plot displays the FSC curve evaluating the overall agreement between the refined atomic model and the experimental cryo-EM density map. Correlation curves are shown for calculations performed with a mask (orange) and without a mask (blue). The horizontal dashed line represents the standard 0.5 FSC validation criterion. The bottom axis indicates the inverse resolution (1/Å), while the top axis shows the corresponding spatial resolution in Å.

**Table S1.** Bacterial strains used in this study.

| Species | Strain | Description | Reference |
| --- | --- | --- | --- |
| *E. coli* | 10β | Competent strain used for plasmid cloning. | New England Biolabs |
| *E. coli* | BL21(DE3) | Competent strain used for recombinant protein expression. | New England Biolabs |
| *E. coli* | BL21(DE3)_SCP H_FIVAR3 | Strain transformed with the pET20b(+)_FIVAR3 plasmid for the production of the recombinant FIVAR3 protein (with N-terminal Strep-Tag II). | This study |
| *E. coli* | BL21(DE3)_SCP H_FIVAR3&ND | Strain transformed with the pET20b(+)_FIVAR3&ND plasmid for the production of the recombinant FIVAR3&ND protein (with N-terminal Strep-Tag II). | This study |
| *S. sanguinis* | 2908 | Wild-type background strain used to generate mutant strains. | Gurung *et al.*, 2016 |
| *S. sanguinis* | SS56 | Reference strain used for the purification of native ScpH filaments. | Mom *et al.*, 2024 |
| *S. sanguinis* | ScpH-mChartreuse | Intermediate mutant strain expressing a ScpH-mChartreuse fusion (generated via sPCR). | This study |
| *S. sanguinis* | ScpH-ALFA-tag | Mutant strain expressing a ScpH-ALFA-tag fusion, used for immunofluorescence microscopy experiments. | This study |

**Table S2.**
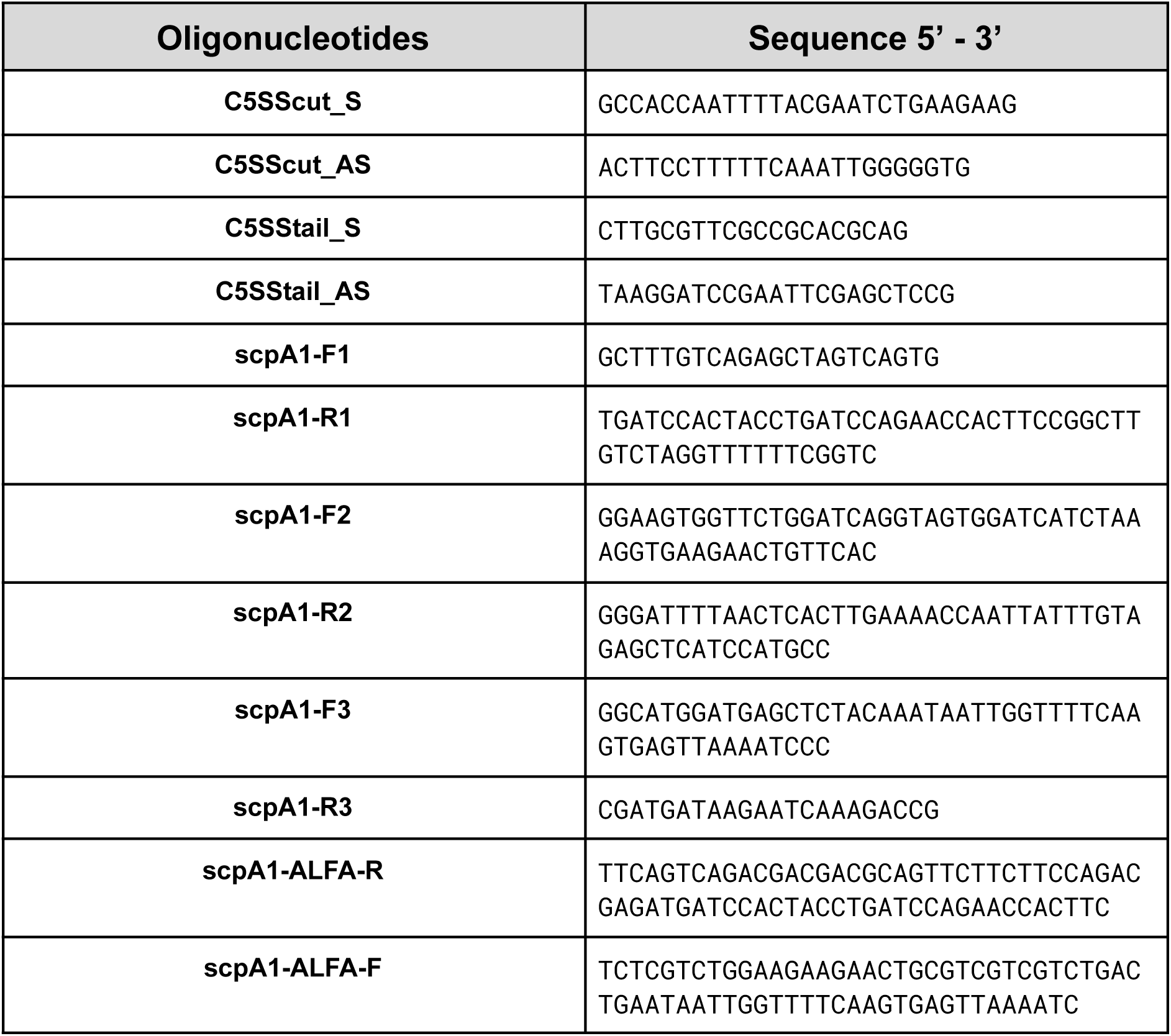
Oligonucleotides used in this study.

**Table S3.** Plasmids used in this study.

| Plasmid | Description | Reference |
| --- | --- | --- |
| prtS_antenna-pET20b(+) | Plasmid containing the C-terminal sequence of ScpH (residues 1197-1506). Used as a template for the construction of pET20b(+)_FIVAR3&ND. | Genscript |
| pET20b(+)_FIVAR3&ND | Vector derived from prtS_antenna-pET20b(+) encoding ScpH FIVAR3 and ND domains with an N-terminal Strep-Tag II. Used for protein expression in *E. coli* BL21(DE3) and as a template to construct pET20b(+)_FIVAR3. | This study |
| pET20b(+)_FIVAR3 | Vector derived from pET20b(+)_FIVAR3&ND encoding only the ScpH FIVAR3 domain with an N-terminal Strep-Tag II. Used for protein expression in *E. coli* BL21(DE3). | This study |
| pNF02-mChartreuse | Plasmid encoding the monomeric green fluorescent protein mChartreuse. Used as a template for the construction of the *S. sanguinis* ScpH-mChartreuse strain. | Fraikin *et al.*, 2025 |

**Table S4.** Cryo-EM data collection and refinement statistics.

| Accession codes |  |
| --- | --- |
| PDB code | 31CH |
| EMDB code | EMD-58274 |
| Data collection |  |
| Microscope | Glacios 2 (Thermo Fisher Scientific) |
| Data collection software | Smart EPU |
| Voltage (kV) | 200 |
| Camera | Falcon 4i (Thermo Fisher Scientific) |
| Energy filter | SelectrisX (Thermo Fisher Scientific) |
| Magnification | 130,000x |
| Electron dose per frame (e<sup>-</sup>/Å<sup>2</sup>) | 1.72 |
| Total electron dose (e<sup>-</sup>/Å<sup>2</sup>) | 51.80 |
| Pixel size (Å) | 0.885 |
| Defocus range (μm) | -0.2 to -2.4 |
| Micrographs acquired (no.) | 25,079 |
| Method | Single particle analysis |
| Processing |  |
| Software used | CryoSPARC |
| Micrographs (no.) | 22,037 |
| Initial particles (no.) | 202,815 |
| Consensus particles (C1) (no.) | 65,221 |
| Symmetry used for local refinement | C2 |
| Final particles after symmetry expansion (no.) | 130,442 |
| Map resolution, masked (Å,FSC 0.143 criterion) | 3.09 |
| Map resolution, unmasked (Å,FSC | 4.3 |
| 0.143 criterion) |  |
| Map sharpening B-factor (Å<sup>2</sup>) | 61.3 |
| **Model refinement and validation** |  |
| Softwares used | ChimeraX, ISOLDE, PHENIX |
| Model-to-map resolution (Å, FSC 0.5 criterion) | 3.40 |
| Chains (no.) | 5 |
| Atoms (no.) | 62,015 |
| Non-hydrogen atoms (no.) | 31,332 |
| Protein residues (no.) | 4,037 |
| R.m.s. deviations - Bond lengths (Å) | 0.31 |
| R.m.s. deviations - Bond angles (°) | 0.48 |
| Clashscore | 1.0 |
| Ramachandran - Favored (%) | 98.0 |
| Ramachandran - Allowed (%) | 2.0 |
| Ramachandran - Outliers (%) | 0.0 |
| Rotamer outliers (%) | 0.0 |
| Atom inclusion in map (%) | 85.1 |
| Q-score | 0.508 |

**Table S5.**
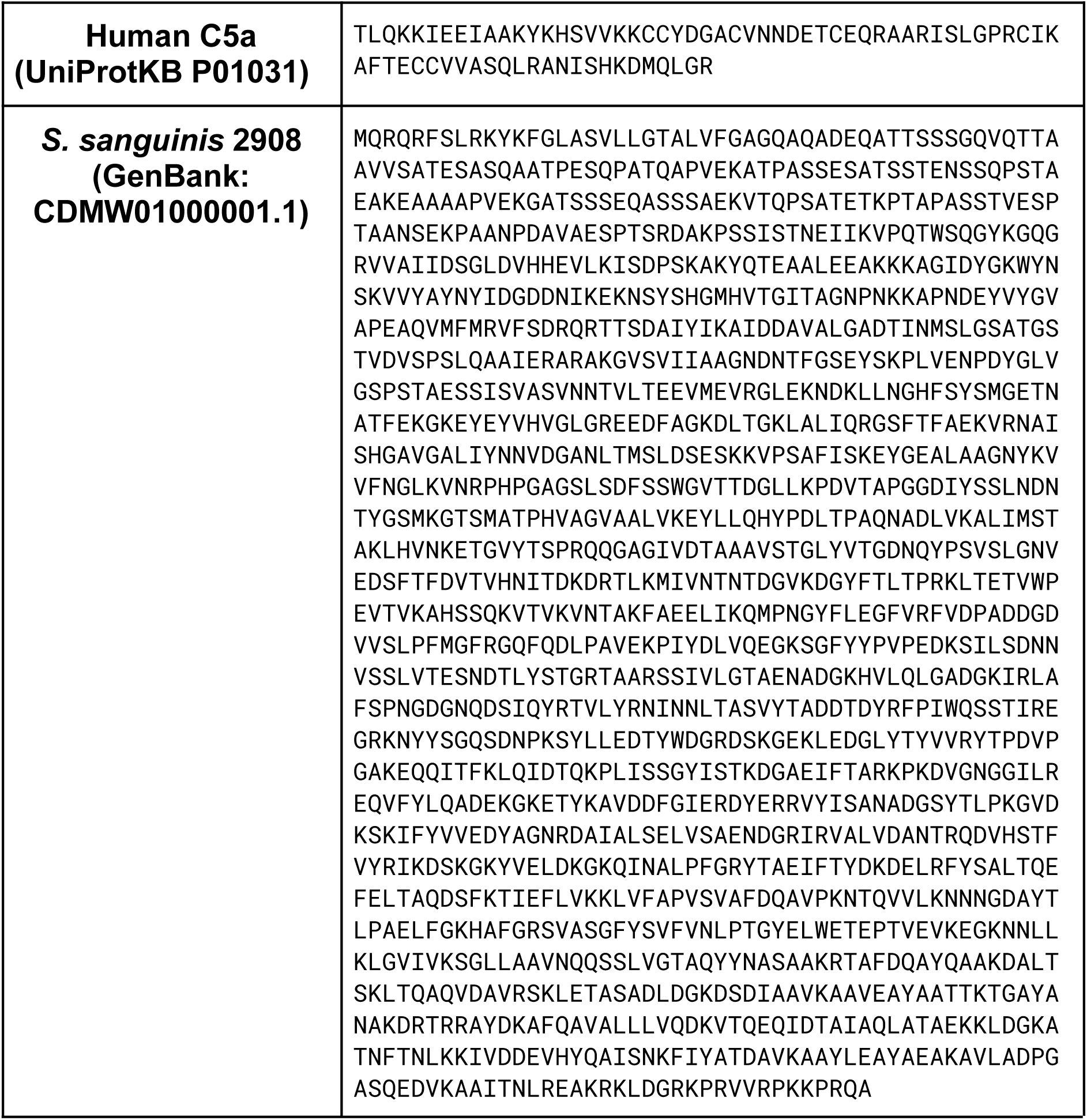
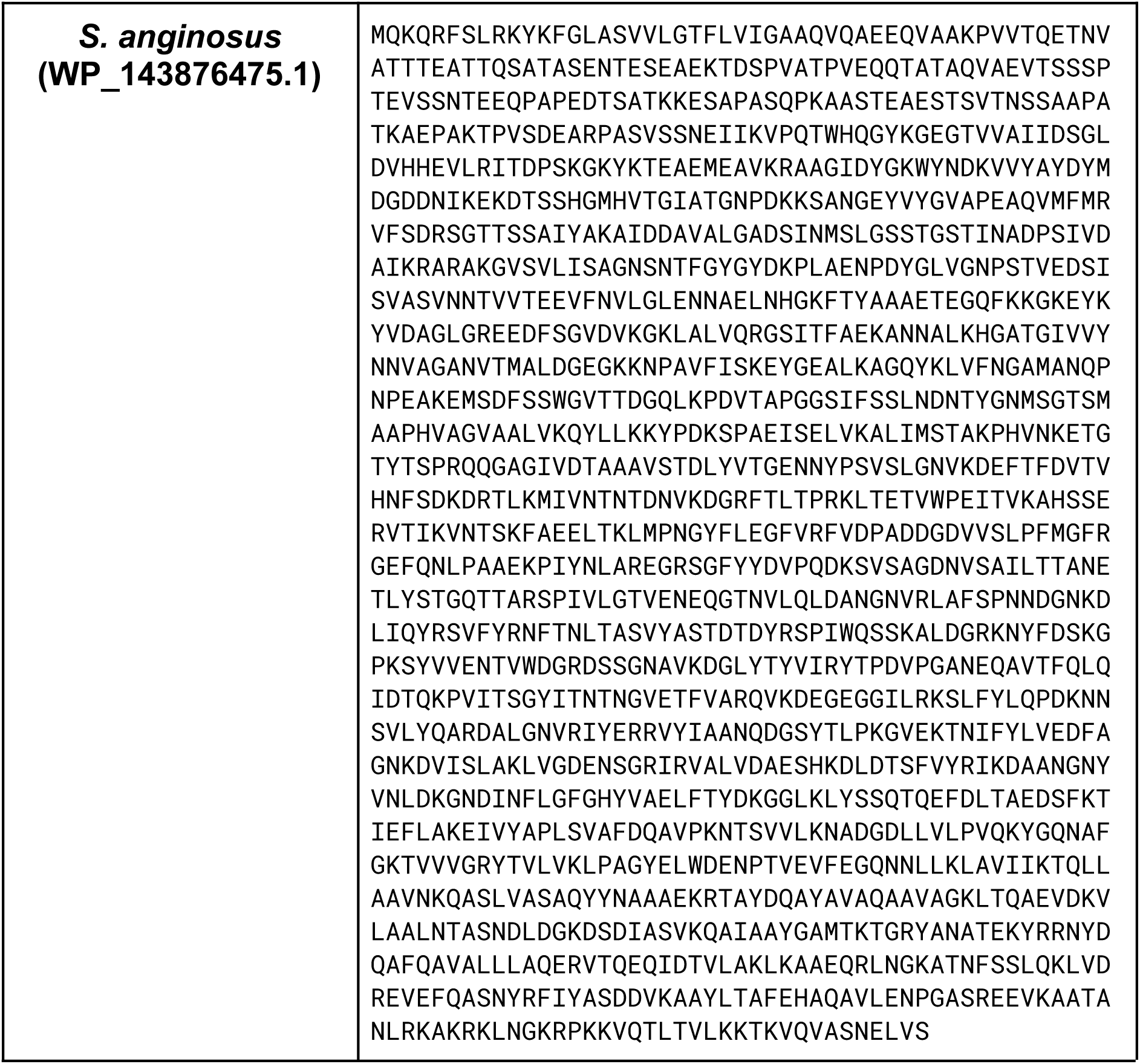
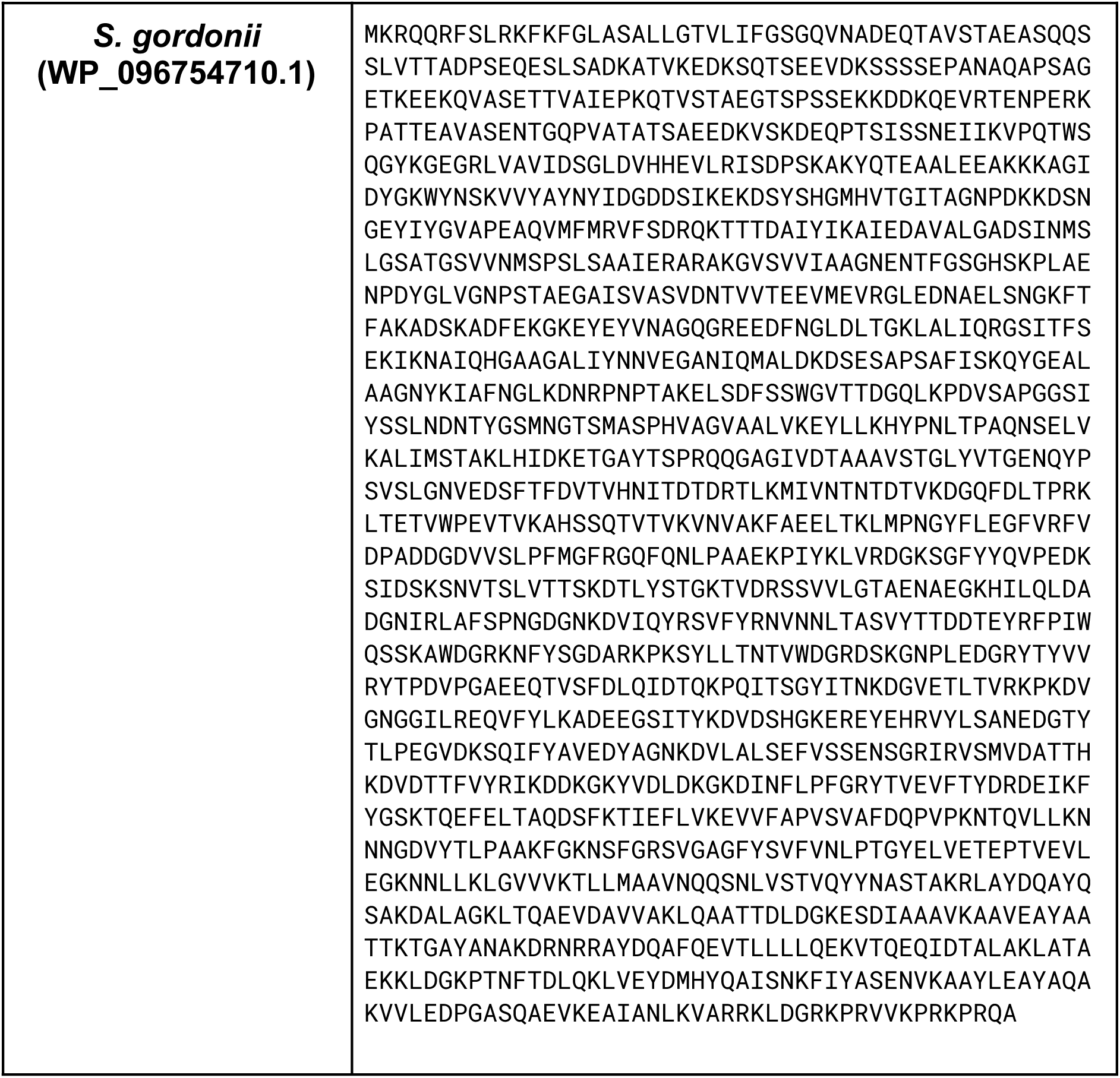
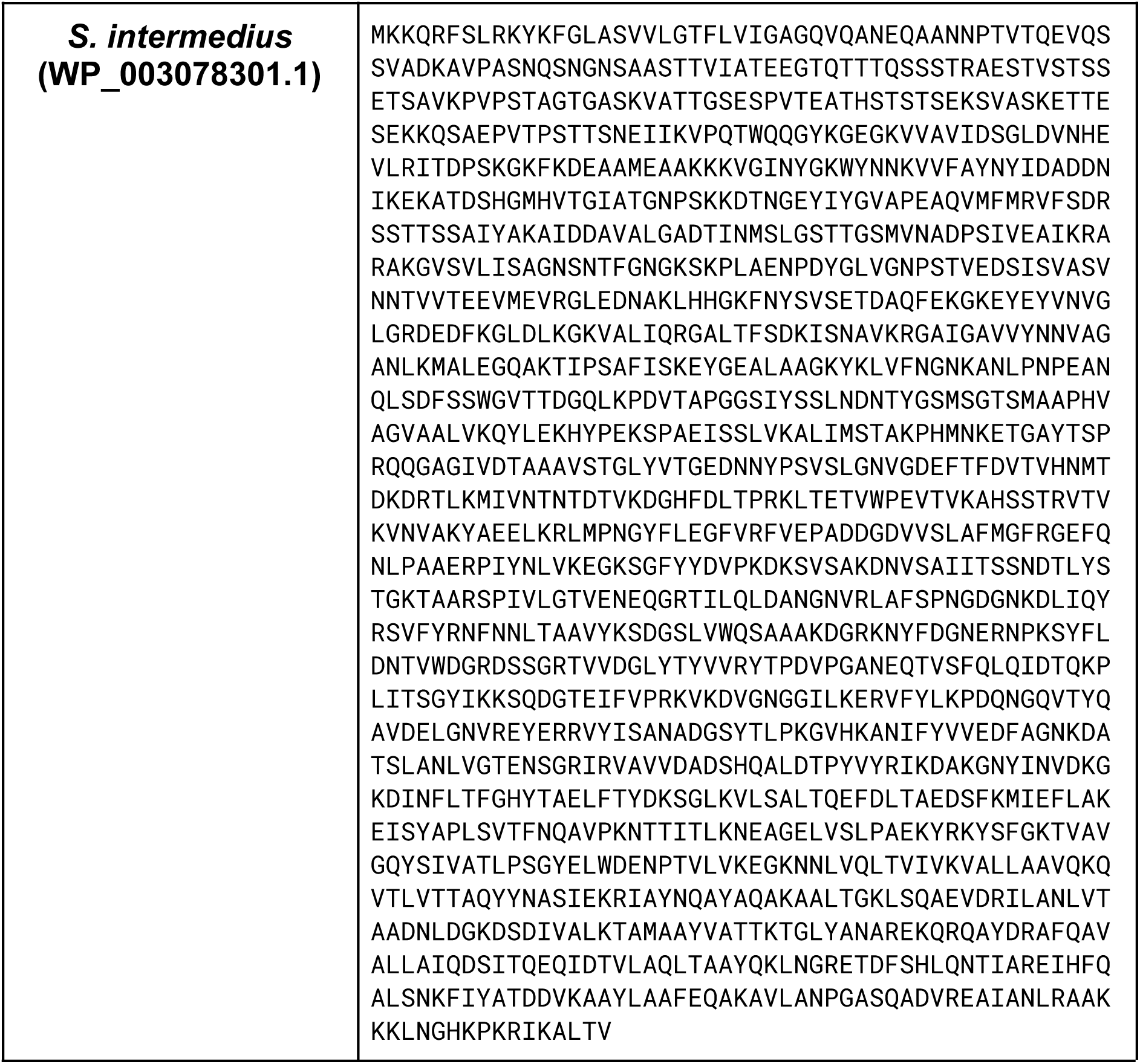
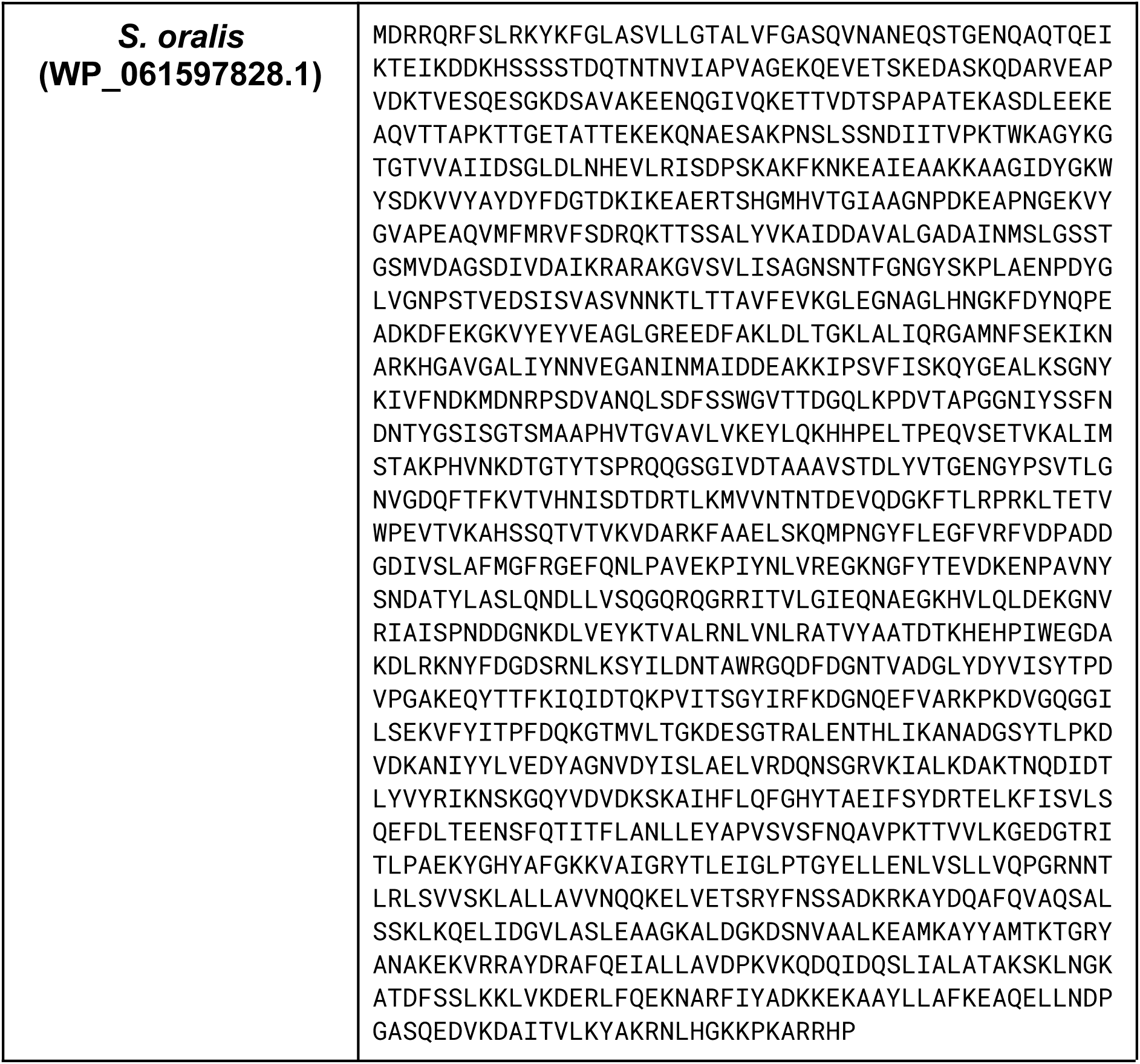
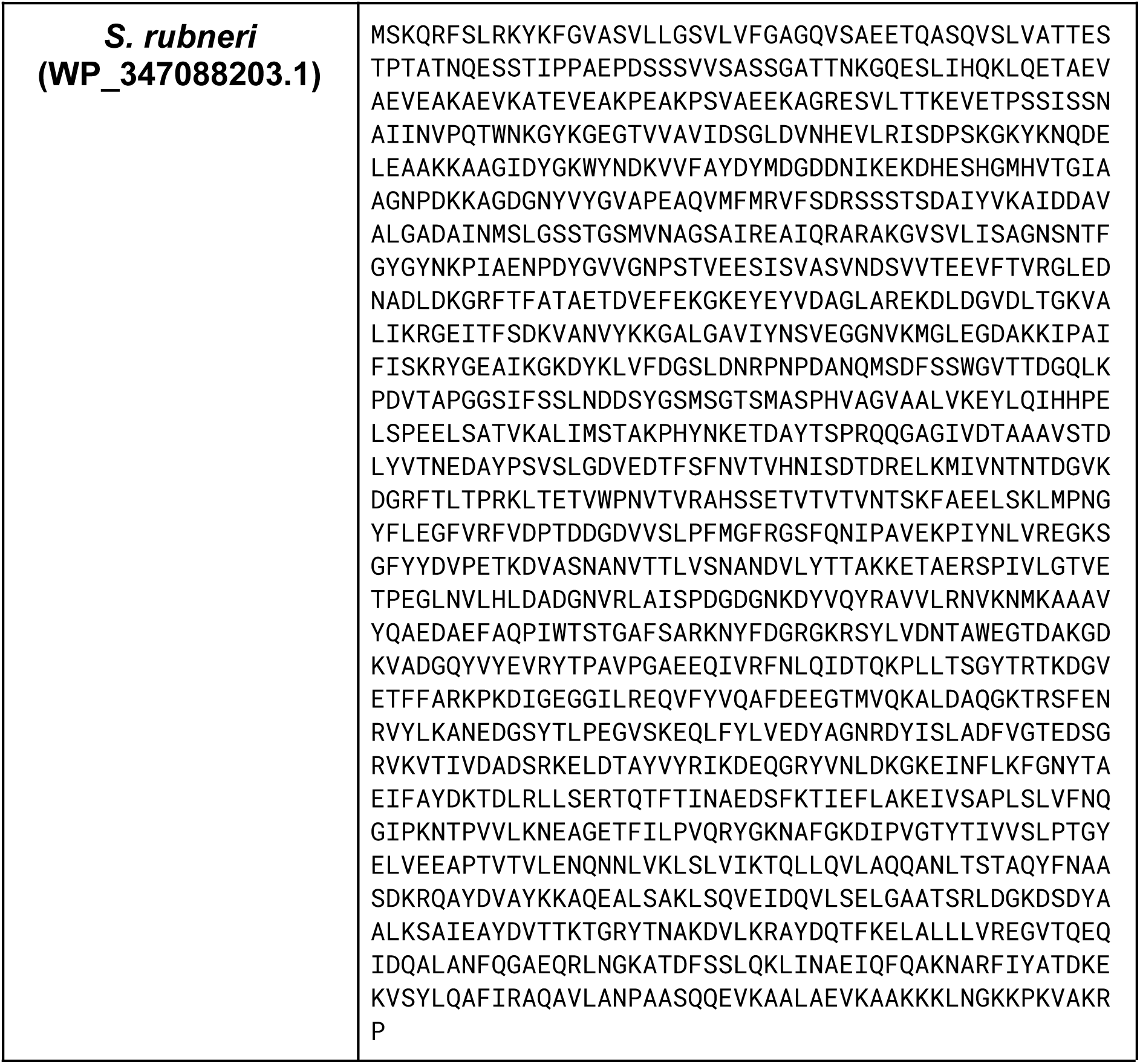
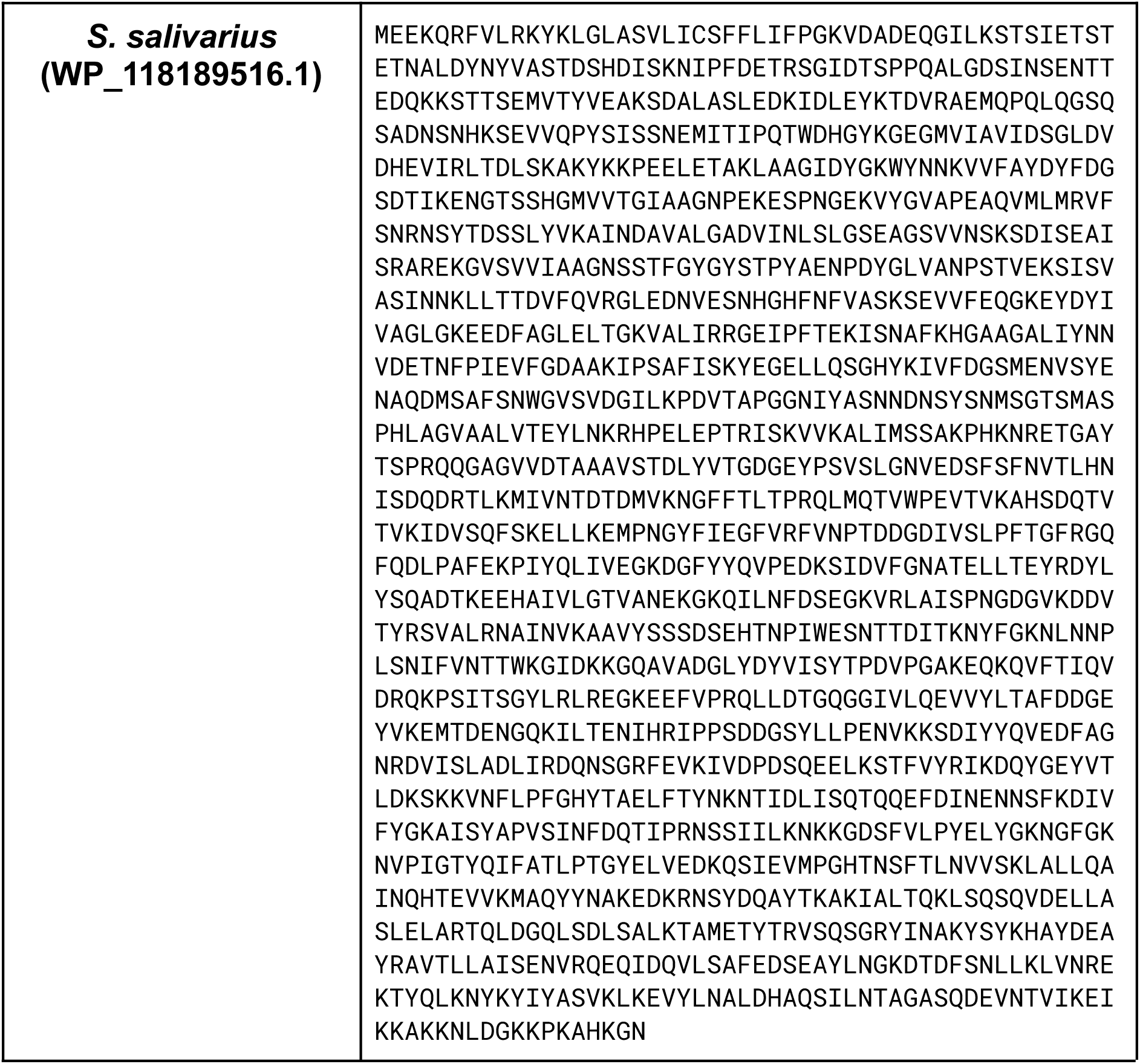
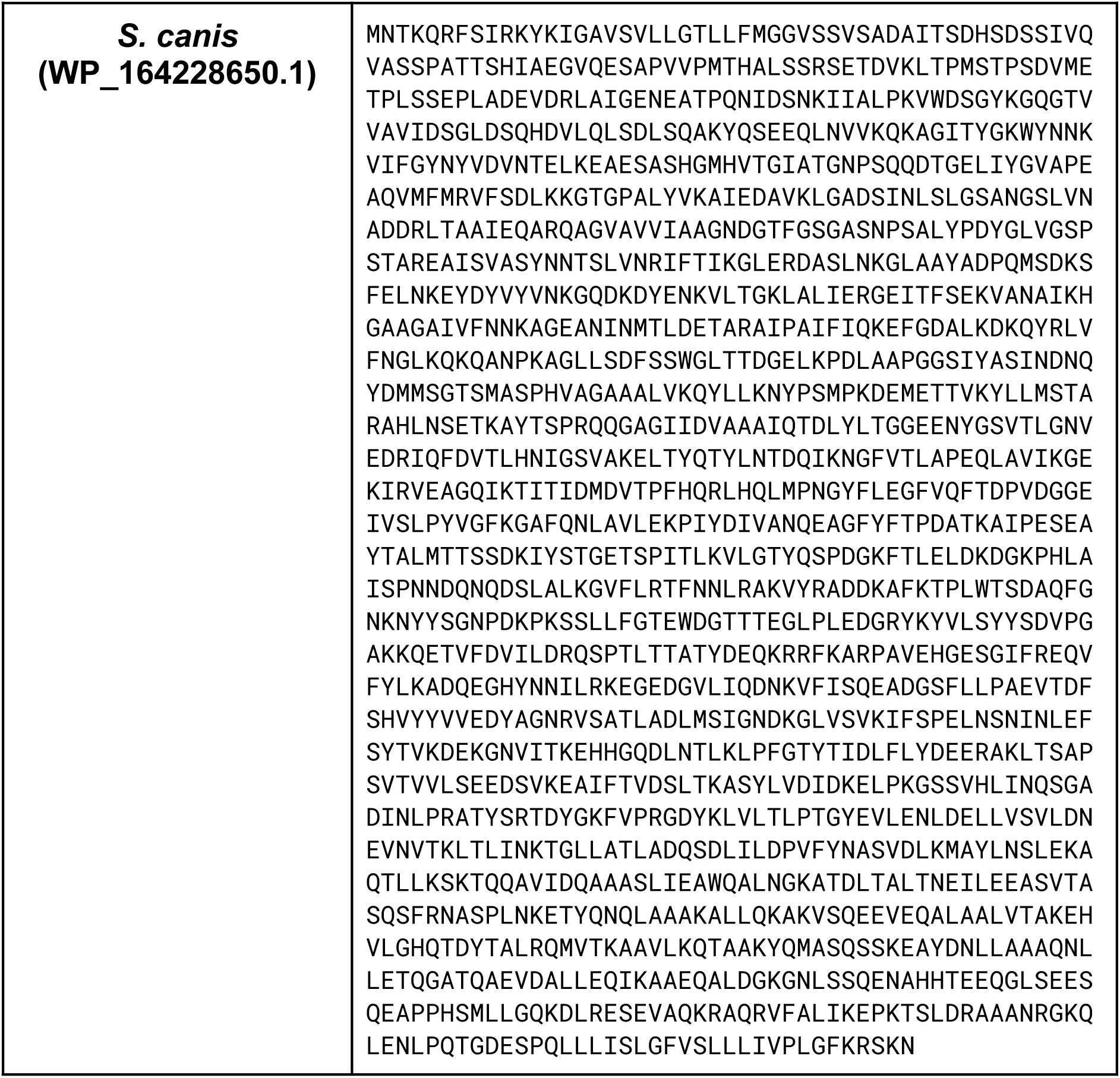
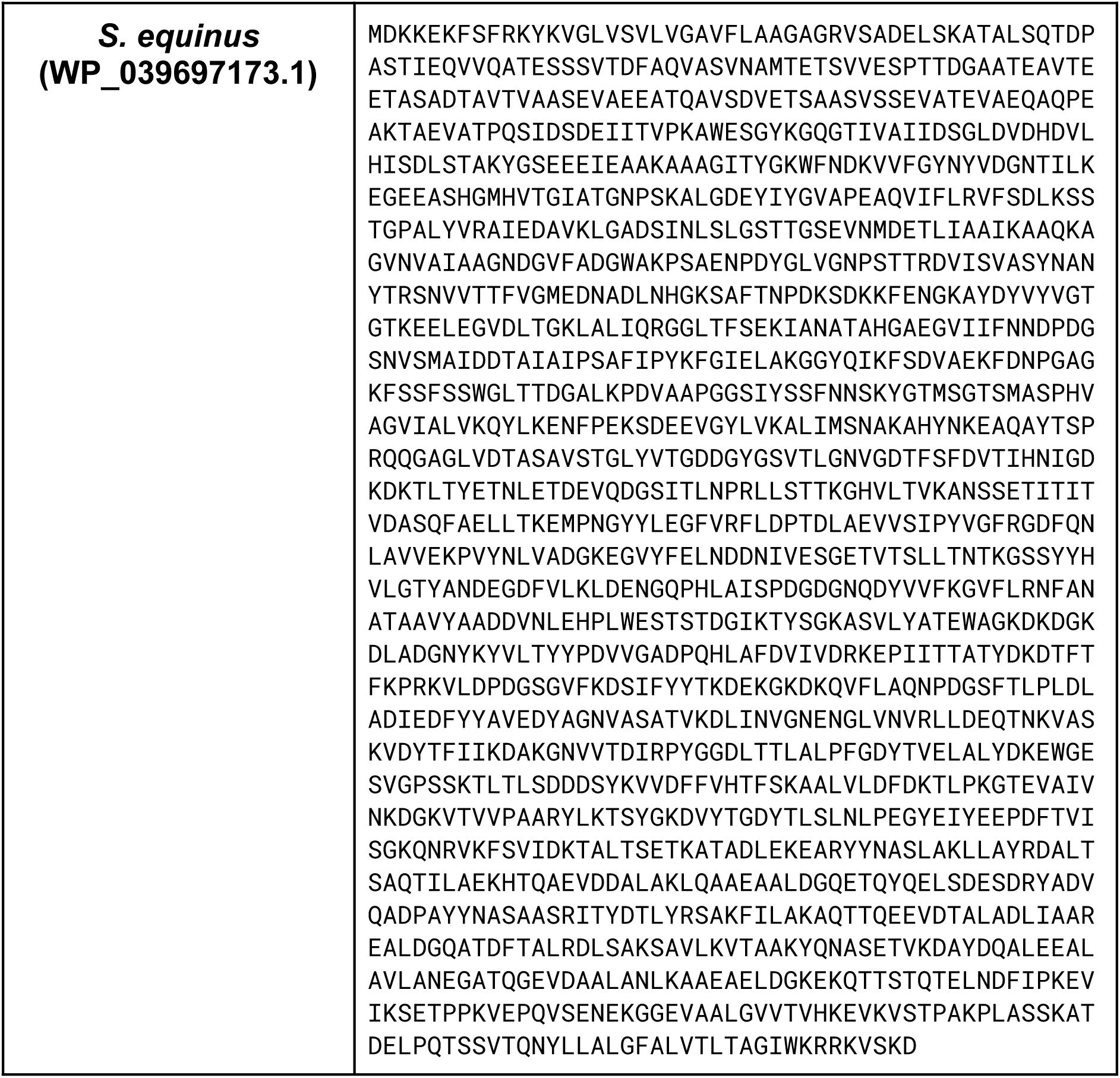
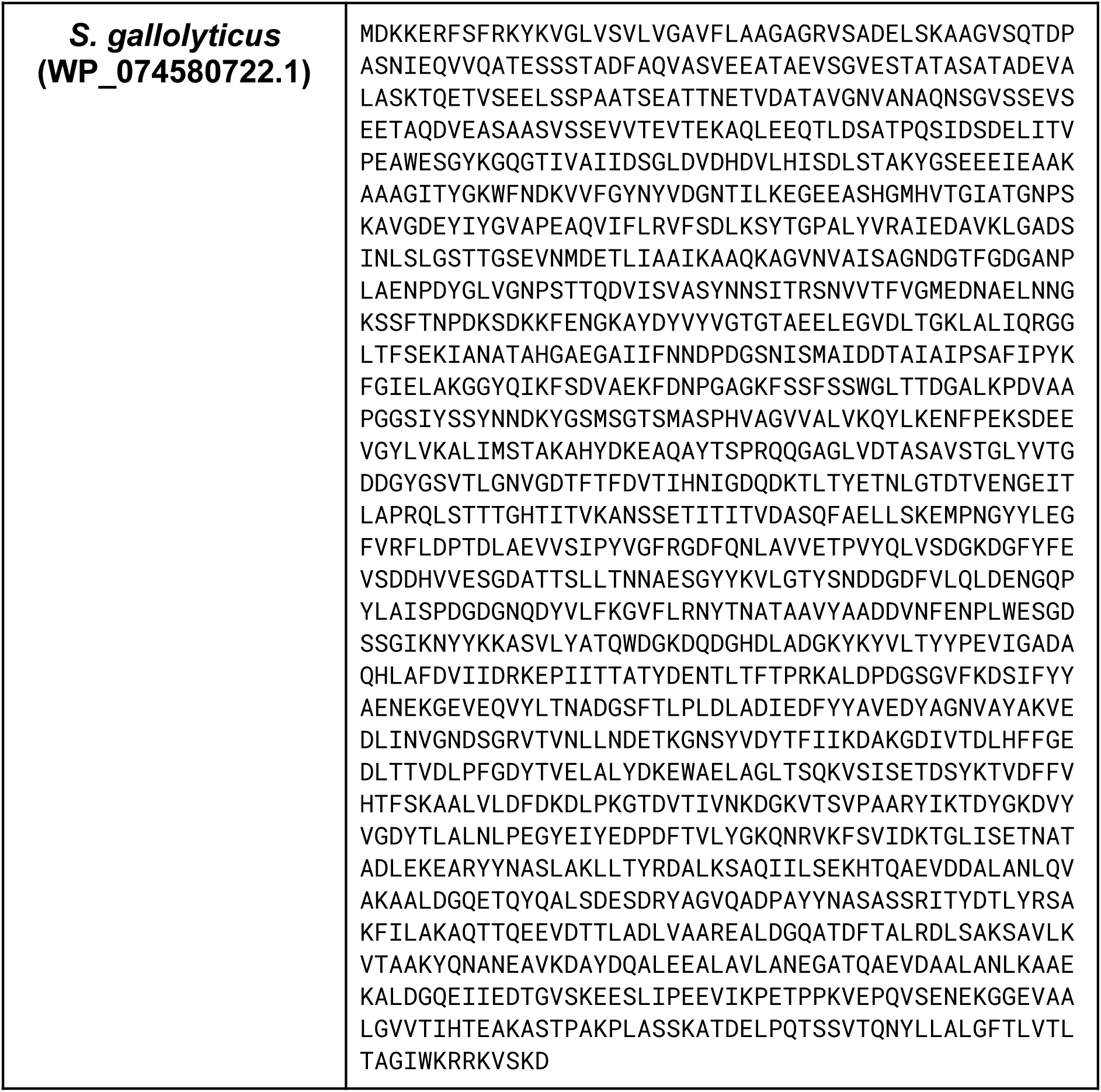
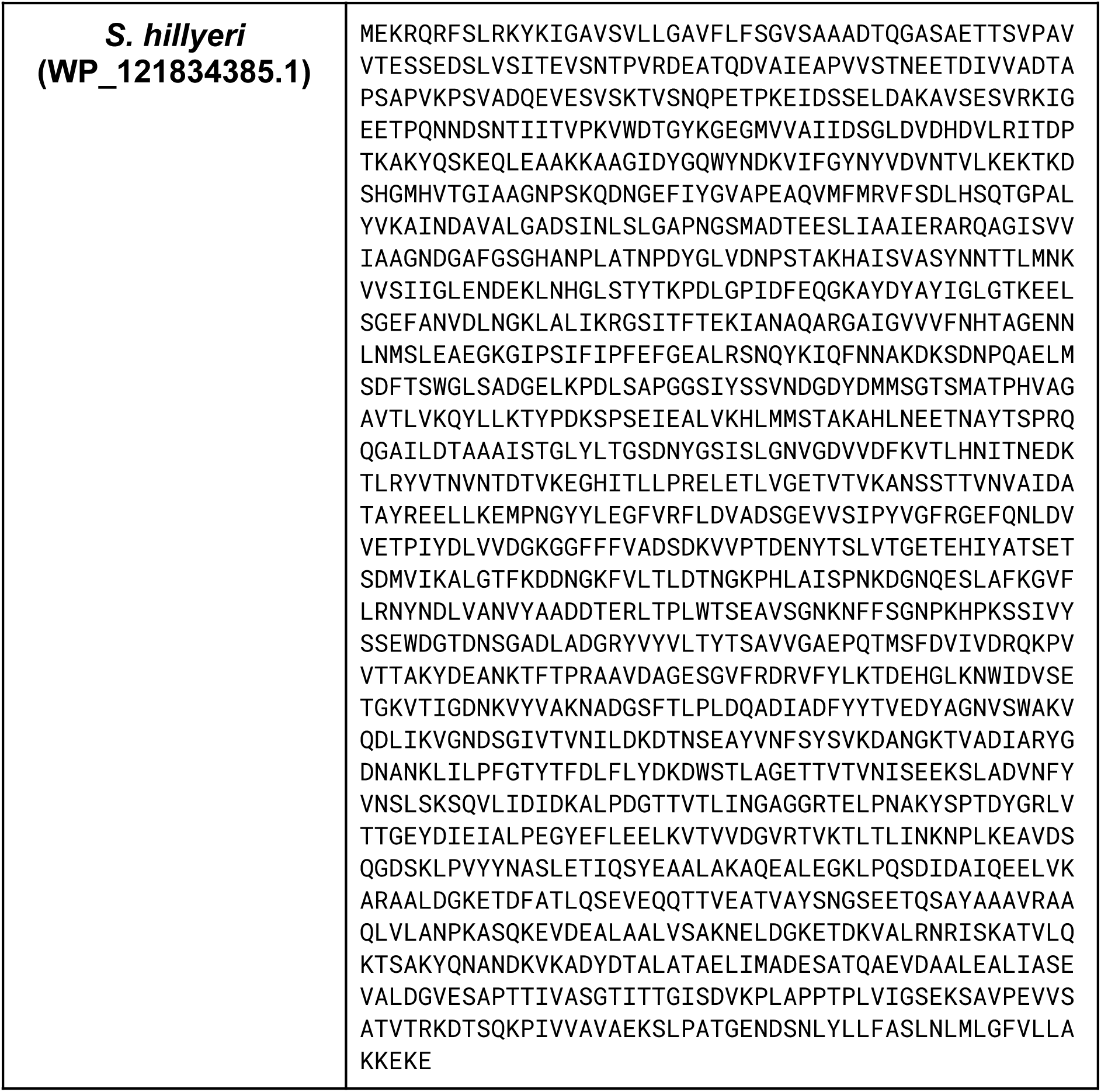
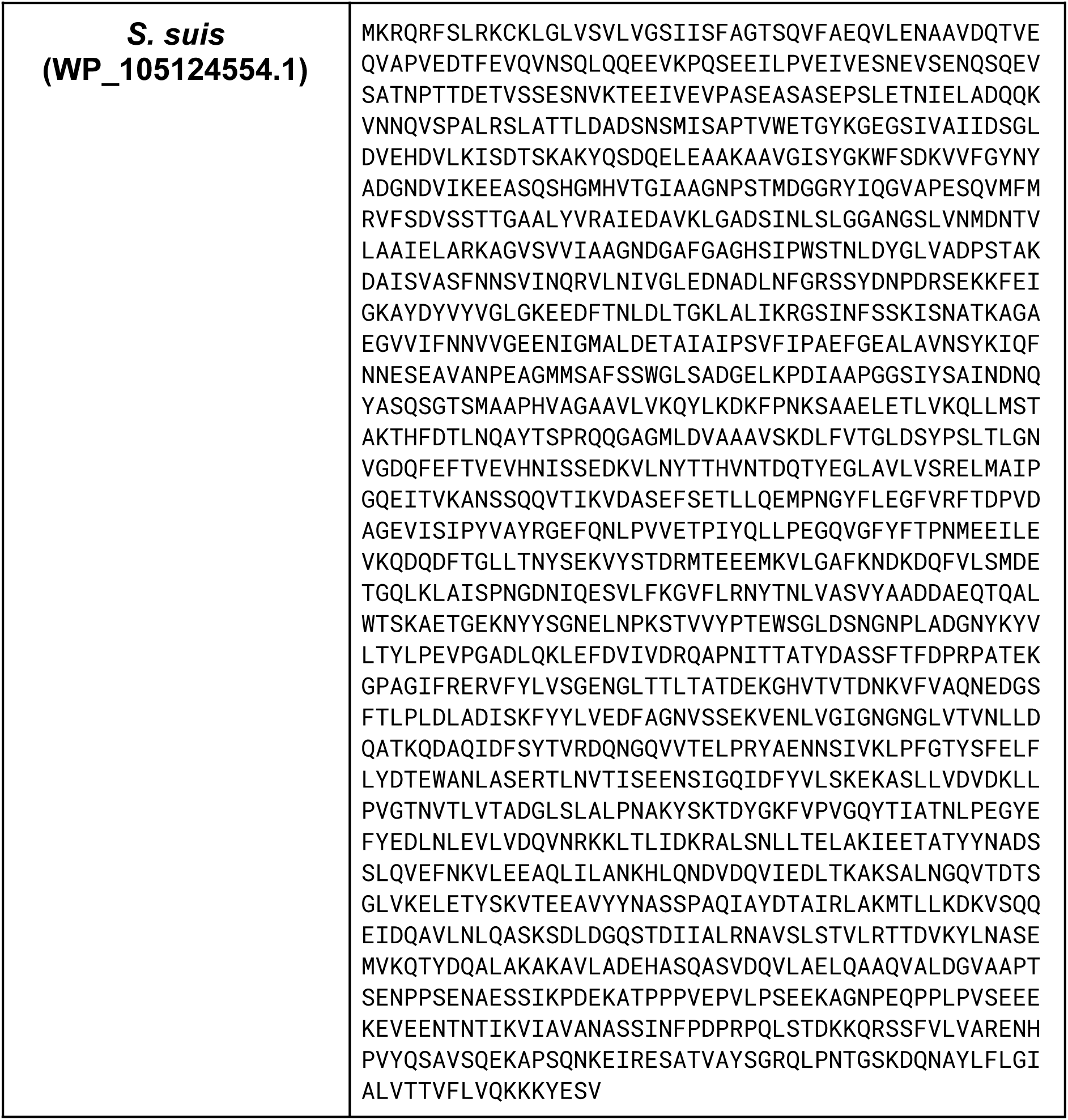
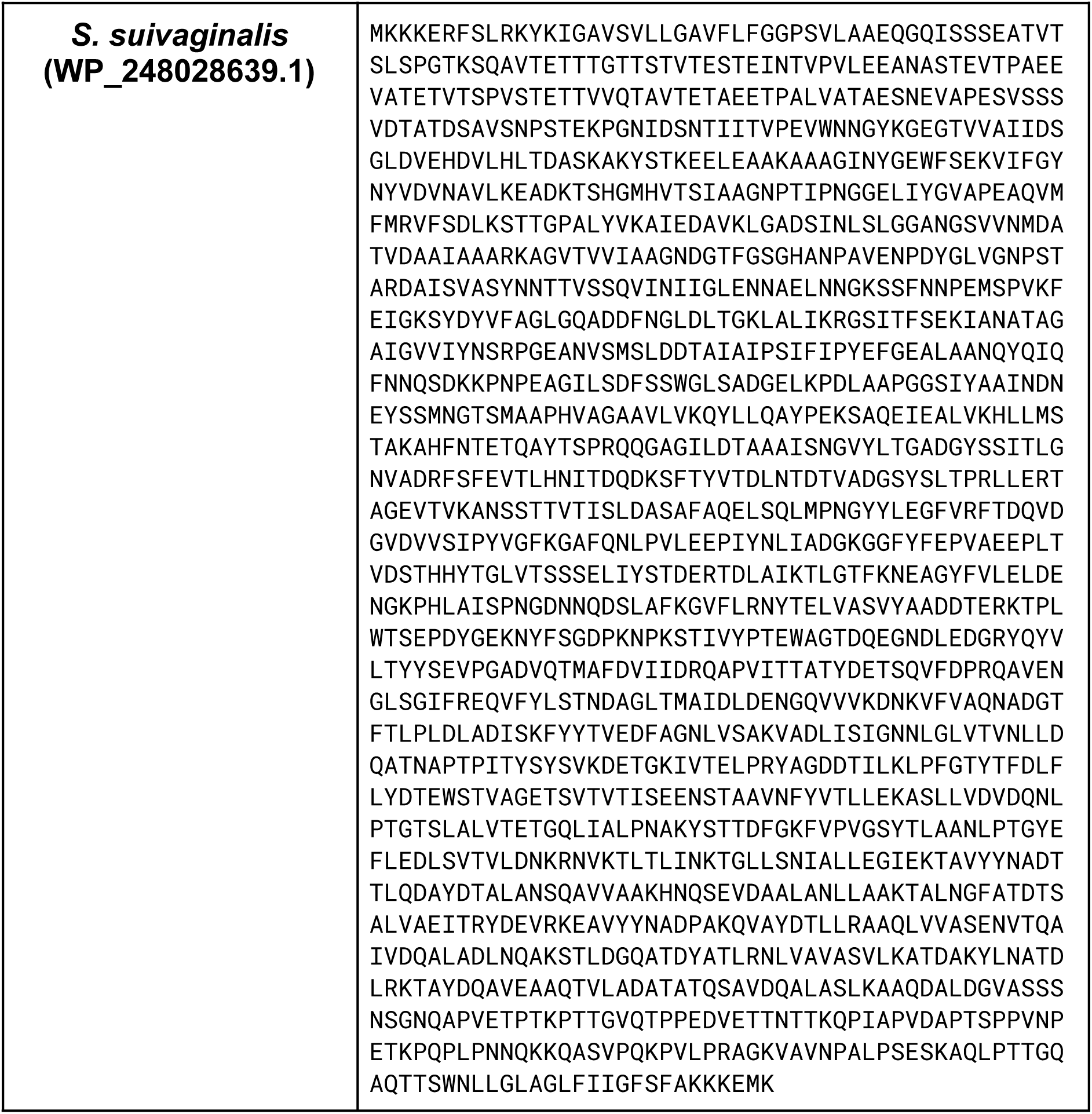
Amino acid sequences used for AlphaFold 3 structural predictions.

